# Somatosensory rhythms and cerebellar-basal ganglia beta-band dynamics in Parkinson’s disease

**DOI:** 10.1101/2025.05.15.653735

**Authors:** Victor Pando-Naude, Lau Møller Andersen

**Affiliations:** Department of Clinical Medicine, Center for Music in the Brain, Aarhus University and The Royal Academy of Music Aarhus/Aalborg, Universitetsbyen, 3-0-17, 8000, Aarhus C, Denmark; Center of Functionally Integrative Neuroscience (CFIN), Aarhus University, Universitetsbyen 3, Building 1710, 8000 Aarhus C, Denmark; Department for Linguistics, Cognitive Science and Semiotics, Aarhus University, Jens Chr. Skous Vej 2, Building 1485, 8000 Aarhus C, Denmark

**Keywords:** Cerebellum, Basal Ganglia, Somatosensory, Timing, Parkinson’s Disease, Beta-Band

## Abstract

Rhythmic cognition relies on the interplay between endogenous and exogenous temporal structures, allowing organisms to anticipate and adapt to events in time. Neural oscillations, particularly in the beta-band (14-30 Hz), have been implicated in this predictive capacity, coordinating sensorimotor networks that support timing and rhythm perception. Parkinson’s disease (PD), a movement disorder characterized by disrupted basal ganglia function, offers a unique framework to probe the neural mechanisms underlying rhythmic cognition. We estimated beta-band source activity and connectivity across cerebellum, basal ganglia, thalamus, and motor cortex during a rhythmic somatosensory timing task to compare brain activity between participants with PD (n=24) and age-matched controls (n=25). Using magnetoencephalography, participants received tactile stimuli in regular (non-jittered) and irregular (jittered) sequences, with the final stimulus omitted to probe prediction *before* and evaluation *after* the omission. We found prediction-related differences between participants with PD and controls in cerebellum and basal ganglia *before* the omissions, while replicating earlier findings in controls, consistent with cerebellar evaluation responses *after* the omissions. In PD, basal ganglia activity showed greater activity for jittered over non-jittered somatosensory rhythms, opposite to controls. Altered cerebellar beta-band responses correlated with PD symptom severity. Connectivity analyses showed group-by-regularity interactions in a sensory-integration network. These findings suggest that temporal prediction and evaluation in rhythmic cognition rely on coordinated beta-band dynamics across the cerebello-striatal-thalamo- cortical network. In PD, disrupted basal ganglia function may be associated with increased cerebellar engagement and altered network dynamics under irregular temporal conditions, potentially reflecting differences in the processing of rhythmic structure.

## 1. Introduction

Rhythms—whether externally perceived or internally generated—allow organisms to predict future states of the environment. This capacity is evident in everyday behaviors such as walking in synchrony with others, anticipating the beat of music, or predicting the timing of speech during a conversation. Across modalities – visual, auditory, and somatosensory – predictive mechanisms support fundamental cognitive functions such as perception, attention, and action by enabling the brain to anticipate and adapt to upcoming events in time ^1,2^. Rhythmic cognition refers to the brain’s capacity to couple endogenous neural oscillations with exogenous rhythms, forming predictive models that optimize sensorimotor behavior. Central to this process are prediction–evaluation cycles, whereby temporal expectations are generated, monitored, and updated when incoming sensory information confirms or violates those expectations. Neural oscillations, particularly in the beta band (14–30 Hz), have been identified as a key mechanism supporting these predictive dynamics across sensory and motor domains ^3^.

In the present study, we distinguish between temporal prediction and temporal evaluation. Prediction-related activity refers to neural processes occurring *before* an expected sensory event and reflecting the anticipation of forthcoming input. Evaluation-related activity refers to neural processes occurring *after* an expected event has been omitted and reflecting the comparison between predicted and actual sensory input.

Two non-cerebral structures, the cerebellum and the basal ganglia, appear to play important roles in rhythmic cognition. The cerebellum has long been implicated in generating internal forward models of sensory feedback ^4,5^, particularly in the somatosensory domain ^6^, allowing precise temporal prediction and evaluation of incoming signal ^7,8^; while the basal ganglia have been associated with interval timing, beat-based processing, and entrainment to regular rhythms, interacting with cortical motor areas to refine temporal predictions and support rhythmic behaviors ^9–11^. Although the present study does not involve beat perception in the classical auditory sense, temporally regular somatosensory stimulation may nevertheless engage predictive timing mechanisms sensitive to rhythmic regularity ^12–16^.

Studies using magnetoencephalography (MEG) show that the cerebellum responds not only to tactile stimuli but also to the omission of expected stimuli, supporting its involvement in maintaining temporal predictions ^12,16^. While debates persist about the differential roles of the basal ganglia and the cerebellum in timing operations, a theory proposes that the basal ganglia are responsible for supra-second timing while the cerebellum manages sub-second timing ^14^. Alternatively, the basal ganglia may handle regular time intervals while the cerebellum processes irregular time intervals ^15,17^. In Andersen & Dalal (2024b) ^18^, we proposed an alternative framework in which the basal ganglia contribute to core timing computations, while the cerebellum integrates temporal information with spatial and sensory cues to generate spatiotemporal predictions. Our model proposes that the cerebellum may not be directly involved in timing *per se*, but instead serves as an integrative hub within a broader basal ganglia-cerebellum-thalamus-M1 system ^18^. This integrated network plays a fundamental role in *predicting* when, where, and what sensory stimuli will occur. From this it follows that timing capabilities may thus be lost due to incapacitation of *either* basal ganglia *or* cerebellum, as also shown by a recent lesion study ^19^. Thus, one might conjecture that neurodegenerative diseases associated with loss of motor control and rhythmicity, such as Parkinson’s disease (PD), would be associated with dysfunction in prediction mechanisms due to altered cerebellar-basal ganglia interactions ^20,21^.

We previously showed that cerebellar beta-band activity following omitted tactile stimuli is modulated by the regularity of preceding input, with higher activity following non-jittered than jittered rhythms, consistent with temporal evaluation of predictable sensory events ^12^. This activity correlates with detection accuracy ^22^, suggesting that cerebellar dysfunction may impair predictive mechanisms critical for smooth movement in PD ^23–26^. PD is primarily characterized by dopaminergic degeneration in the substantia nigra pars compacta, disrupting basal ganglia-thalamo-cortical circuitry and impairing motor functions ^27,28^, including timing ^29^. By comparing participants with PD to age-matched healthy controls, we can gain insights into the fundamental mechanisms of (1) sensory processing, (2) temporal prediction and evaluation, and (3) cerebellar beta-band activity ^30^. At the same time, this comparison can help reveal potential cerebellar dysfunction specifically linked to PD. To meet these ends, we used MEG.

Due to the high temporal resolution of MEG, it is possible to distinguish effects that happen *before* the expected but omitted stimulus, i.e. related to the *prediction* of upcoming stimuli, from effects that happen *after* the expected but omitted stimulus, i.e. related to the *evaluation* of predictions. MEG captures millisecond-scale neural dynamics, complementing the high spatial but limited temporal resolution of magnetic resonance imaging. MEG has traditionally been regarded as mainly sensitive to the cerebral cortex, but recently it has been shown that the cerebellum is within reach of MEG ^31,32^. The sensitivity of MEG to the basal ganglia, on the other hand, is low, especially in situations where the cerebral cortex is activated simultaneously ^33^. However, accumulating evidence suggests that subcortical contributions are detectable under appropriate conditions. Computational modeling studies have shown that the basal ganglia can generate measurable magnetic fields ^33,34^, and empirical findings suggest basal ganglia involvement in timing tasks captured with MEG ^24,35^. Importantly, omission paradigms reduce concurrent cortical activity, thereby increasing sensitivity to subcortical signals. In other words, because no physical stimulus is delivered at the expected time point, omission paradigms avoid the large evoked cortical responses typically elicited by sensory stimulation, potentially improving sensitivity to neural activity originating from deeper structures. These advances support the possibility that MEG source reconstruction may estimate source-level activity consistent with basal ganglia involvement in the current paradigm ^34,36^.

Despite extensive work on temporal processing, it remains unclear whether predictive timing depends primarily on computations within the basal ganglia, the cerebellum, or coordinated interactions between these structures. Omission paradigms provide a useful framework for addressing this question because they allow prediction-related activity before an expected event to be dissociated from evaluation-related activity following an omitted event. Beta-band oscillations are particularly relevant because they have consistently been implicated in temporal prediction and sensorimotor timing ^23,37^. PD provides a unique opportunity to investigate these mechanisms because basal ganglia dysfunction may reveal how predictive timing depends on cerebello–basal ganglia dynamics.

This study aims to investigate beta-band dynamics associated with rhythmic temporal prediction in PD, focusing on cerebellar and basal ganglia involvement during somatosensory rhythmic omissions. Using MEG, we estimated beta-band source activity in response to regular (non-jittered) and irregular (jittered) sequences of tactile stimuli and omissions, comparing participants with PD to age-matched healthy controls. This comparison also provided an opportunity to replicate and assess the robustness of the omission-related cerebellar beta-band effect previously reported in healthy adults (Andersen & Dalal, 2021) ^12^.

The hypotheses of interest were:

1. Based on previous omission studies ^12,18^, control participants were expected to show greater cerebellar beta-band activity in omissions *after* regular (non-jittered) compared with irregular (jittered) somatosensory rhythms, reflecting temporal evaluation of predictable sensory events.
2. Participants with PD, compared to controls, were expected to show distinct beta-band activity between regular (non-jittered) and irregular (jittered) rhythms in relation to the expected but omitted stimulus in cerebellum and basal ganglia, reflecting altered temporal processing.

For exploratory reasons, we also estimated the functional connectivity in the beta-band between the cerebellum, the basal ganglia, the thalamus and the primary motor cortex. We have suggested that these areas together comprise a temporal integration network enabling proactive action ^18^. Within this framework, the basal ganglia are proposed to contribute temporal information relevant for predicting when events will occur, whereas the cerebellum is proposed to integrate this temporal information with sensory and spatial information to generate spatiotemporal predictions. The thalamus is proposed to act as a relay and integration hub, coordinating information exchange between cerebellar, basal ganglia, and cortical motor systems, thereby enabling the implementation of predictive signals in ongoing behavior ^22^.

## 2. Materials and Methods

### Participants

Participants in the study were individuals diagnosed with PD, according to the Movement Disorder Society (MDS) diagnostic criteria ^38^, as well as an age-matched control group without PD, henceforth “controls”. All participants in the study were right-handed. The study was approved by the regional ethics committee (De Videnskabsetiske Komitéer for Region Midtjylland) in accordance with the Declaration of Helsinki. This study was not preregistered.

Participants were recruited following ethical guidelines and provided informed consent before participating in the study. Fifty-six participants were recruited for the study. Of these, three were excluded due to MEG artifacts caused by metallic implants, three due to left-handedness, and one following visual inspection of the structural MRI revealing cerebellar atrophy in consultation with a local radiologist, resulting in a final sample of 49 participants.

No formal *a priori* power analysis was performed. The sample size was determined based on previous MEG studies using comparable somatosensory omission paradigms ^12,39^ and task-based PD designs ^40,41^, which typically include approximately 20–30 participants per group ^39,40^. Earlier studies using the present paradigm reported medium-to-large cerebellar beta-band omission effects in healthy participants ^12,22^. The current study was therefore designed to provide sufficient sensitivity to detect comparable effects while remaining feasible within the constraints of MEG data acquisition in a clinical population.

Detailed demographic information was recorded for each participant (**Table 1**). No data on socioeconomic status, communities of descent, or race/ethnicity was collected. Groups were age-matched at the group level, with mean ages differing by ∼6 years (controls: 60.2 ± 9.4 years; PD: 66.0 ± 7.8 years), but not individually paired by age. Gender/sex was determined by asking the participants. Sex distribution was not matched, with a higher proportion of females in the control group. Participants were screened for mild cognitive impairment using the Montreal Cognitive Assessment (MoCA) ^42^ and depression symptoms using the Major Depression Inventory (MDI) ^43^ and were excluded if they scored less than 26 and higher than 24, respectively. None were excluded based on these scores.

**Table 1.** Demographics.

|  | Controls | PD | U | p |
| --- | --- | --- | --- | --- |
| Participants (n) | 25 | 24 | - | - |
| Sex/gender | F=16/M=9 | F=8/M=16 | $\chi^2=3.46$ | 0.0628 |
| Age (mean/SD years) | 60.2 ± 9.43 | 66.0 ± 7.77 | 415 | 0.0219 |
| MDI (median/range points) | 3 (0-19) | 8 (0-17) | 467 | 0.000831 |
| MoCA (median/range points) | 29 (26-30) | 29 (26-30) | 311 | 0.829 |
| PD dx (median/range years) | - | 4 (1-17) | - | - |
| UPDRS-III (median/range points) | - | 31 (13-47) | - | - |
| Hoehn and Yahr (median/range points) | - | 2 (1-2) | - | - |
| LEDD (median/range milligrams) | - | 470 (235-1014) | - | - |

PD, Parkinson's disease; MDI, Major Depression Inventory; MoCA, Montreal Cognitive Assessment; PD dx, years since PD diagnosis; UPDRS-III, Motor Examination of the Movement Disorder Society Unified Parkinson's Disease Rating Scale; LEDD, levodopa equivalent daily dose in milligrams; U, Mann-Whitney U Test; p, corrected p-value, $\chi^2$ , Chi-squared test.

To assess the severity of PD, the motor examination of the Movement Disorder Society’s Unified Parkinson’s Disease Rating Scale (UPDRS-III) ^44^ was performed by the first author (Victor Pando-Naude, MD, PhD), immediately prior to data acquisition, such that participants were approximately in the same motor state during clinical evaluation and MEG. Participants with PD were instructed to continue their regular dopaminergic medication regimen and were therefore tested in the ON-medication state during the UPDRS-III evaluation and MEG acquisition (**Figure S2, Table S1**). MEG recordings were acquired a median of 90 min after the most recent levodopa dose (mean = 97.9 min, range = 0–240 min). Levodopa equivalent daily dose (LEDD) was calculated for each PD participant based on the study by Tomlinson et al. (2010) ^45^. Control participants did not report use of dopaminergic medication. Motor fluctuations and dyskinesia were not systematically quantified during the study; however, no severe dyskinesia interfering with motor evaluation or MEG acquisition was observed during the study (**Table S2**).

### Stimuli and Procedure

Tactile stimulation was administered using two ring electrodes powered by an electric current generator (Stimulus Current Generator, DeMeTec GmbH, Germany). These ring electrodes were affixed to the tip of the right index finger, with one placed 10 mm below the bottom of the nail and the other 10 mm below that.

Stimuli were delivered using PsychoPy (version 3.2.4) ^46^ on a Linux operating system running Ubuntu Mate (kernel version 4.15.0-126). Six short breaks were administered throughout the session to ensure participant comfort and to verify their well-being through verbal communication with the experimenter.

Stimuli were presented in sequences of six, with an inter-stimulus interval of 1,497 ms to prevent synchronization with the 50 Hz power line interference. Each pulse had a duration of 100 µs, and the level of current was individually adjusted for each participant. In each sequence, the sixth stimulus was followed by an omission, marking the start of a new stimulation train. Thus, a 2,994 ms interval elapsed between the last stimulus of one train and the first stimulus of the next. Three types of sequences were administered (**Figure 1**):

**1.** Non-Jittered Somatosensory Rhythms: All stimuli occurred precisely on time (0% jitter).
**2.** Jittered Somatosensory Rhythms: Stimuli 4–6 were jittered by 15%, occurring from −225 ms to +225 ms relative to the non-jittered sequence. The jitter administered on a given stimulus is drawn from a uniform distribution in steps of 1 ms.
**3.** Non-Stimulation Sequence: A train of seven non-stimulations, each lasting 1,497 ms, occurred after every fifteen stimulation sequences.

**Figure 1.**
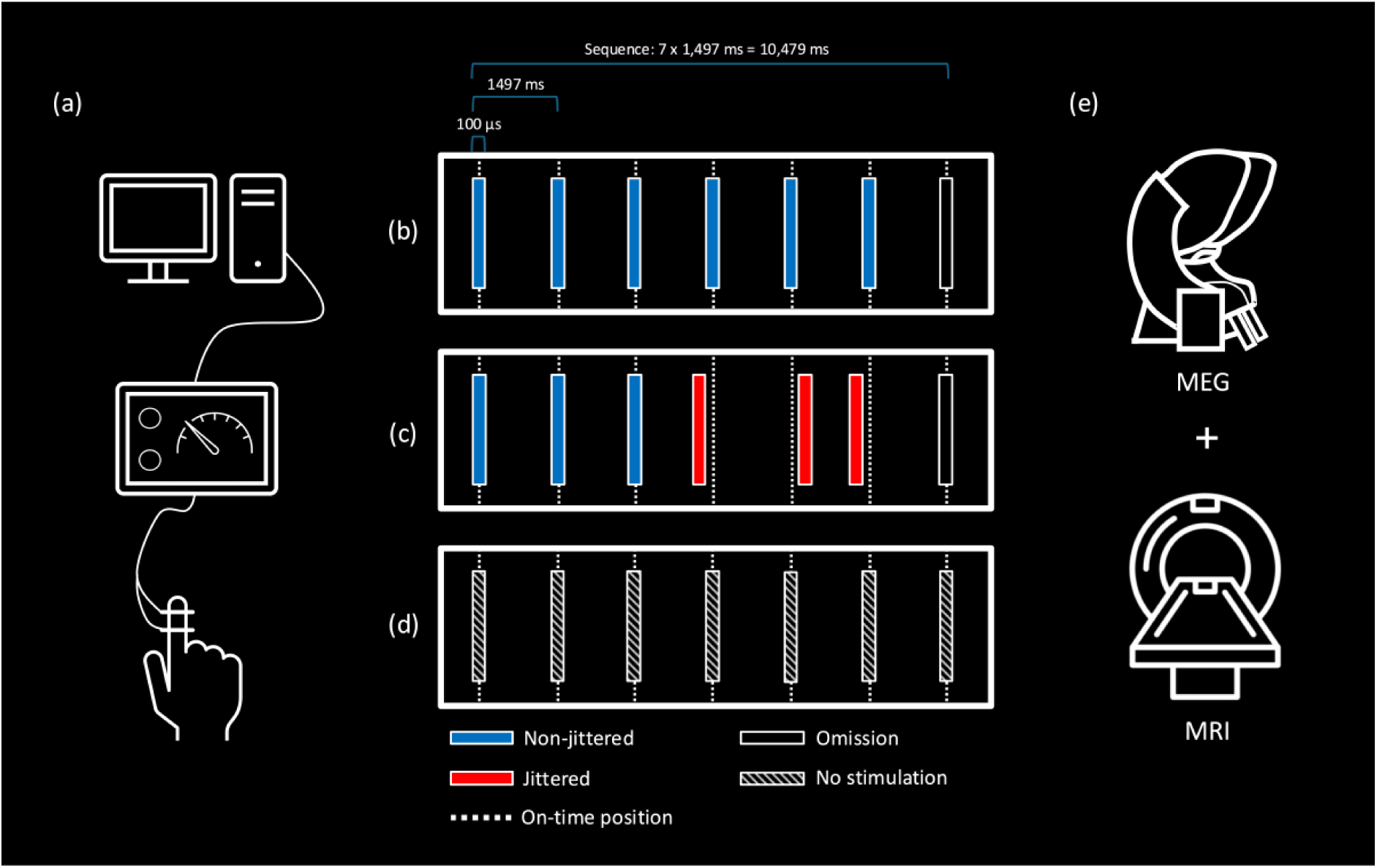
Experimental paradigm. **(a)** Tactile stimuli were administered via ring electrodes placed on the right index finger, delivering sequences of six pulses with a duration of 100 µs and inter-stimulus interval of 1,497ms. PsychoPy controlled the stimulation, while participants watched a nature documentary and remained motionless. EOG, ECG, and EMG recordings were collected to monitor eye movements, heart activity, and neck muscle tension. Stimulus sequences included **(b)** non-jittered somatosensory rhythms, **(c)** jittered somatosensory rhythms (15%), and **(d)** non-stimulation sequences. **(e)** MEG was acquired during the paradigm, and structural MRI was acquired on a separate day within two weeks of the MEG session.

Each stimulation sequence type was administered 150 times, while the non-stimulation sequence was administered 30 times. The sequences were pseudorandomly interleaved and counterbalanced to mitigate order effects. The use of six stimuli per train and 15% temporal jitter was selected a priori based on previous studies using the same somatosensory omission paradigm, in which these parameters reliably elicited omission-related cerebellar beta-band effects ^18,22^. Participants watched a silent nature documentary throughout the experiment to maintain wakefulness and reduce boredom during the recording session. Participants were instructed to attend to the documentary rather than the tactile stimulation. The study was not designed to assess conscious perception of the stimuli. The non-stimulation condition was included as part of the original paradigm design to provide periods without tactile stimulation but was not directly relevant to the present hypotheses concerning temporal prediction and evaluation under regular and irregular stimulation conditions. Therefore, the non-stimulation sequences were not further analyzed. (**Figure 1**, **Table 1**).

### Preparation of Participants

Participants underwent standard MEG preparation, including Electrooculography (EOG), electrocardiography (ECG), electromyography (EMG), and respiration recordings to monitor eye movements, cardiac activity, muscle tension and breathing. EOG electrodes were placed around the eyes, ECG on the collarbones, and EMG on the splenius muscles. Four HPI coils were attached to the head, and head shape digitization was performed using Polhemus FASTRAK, including fiducials and ∼200 extra points. Participants were positioned supine in the MEG scanner with ring electrodes secured to the right index finger. Stimulation current was individually adjusted: starting at 1.0 mA, increased until perceptible but not irritating, and then fine-tuned for comfort.

### Data Acquisition

Data was acquired at Aarhus University Hospital (AUH), Denmark. Whole-head MEG data was acquired using an Elekta Neuromag TRIUX system, comprising 306 sensors arranged as 102 magnetometers and 204 planar gradiometers, located within a magnetically shielded room (Vacuumschmelze Ak3b). The sampling frequency was 1,000 Hz; online filtering was applied to the acquired data, consisting of a low-pass filter with a cutoff frequency of 330 Hz and a high-pass filter with a cutoff frequency of 0.1 Hz. No EEG data was recorded.

### MEG Data Processing

MEG data were processed using MNE-Python ^47^ following Andersen (2018) ^48^. Bad channels (mean = 0.22 per participant) were excluded based on power spectra inspection. For evoked responses, data were low-pass filtered at 40 Hz [one-pass, noncausal, finite impulse response; zero phase; upper transition bandwidth: 10.00 Hz (−6 dB cutoff frequency: 45.00 Hz; filter length 331 samples; a 0.0194 passband ripple and 53 dB stopband attenuation)], epoched (−200 to 600 ms), baseline-corrected (− 100 ms – 0 ms), and averaged by condition. Analyses focused specifically on the beta band (14–30 Hz), based on prior studies implicating beta-band dynamics in temporal prediction and omission processing ^12,39^. Beta-band responses were extracted by band-pass filtering [14–30 Hz; lower transition bandwidth: 3.50 Hz (−6 dB cutoff frequency: 12.25 Hz); upper transition bandwidth: 7.50 Hz (−6 dB cutoff frequency: 33.75 Hz); filter length 943 samples; passband ripple: 0.0194; stopband attenuation: 53 dB], Hilbert transforming, and epoching from −750 to +750 ms around each stimulus or omission (demeaned using the interval from −750 ms to 750 ms). The main analysis of interest was the beta band activity. Evoked responses were extracted to confirm the presence of the expected SI and SII components, in line with previous findings using the same paradigm ^12^ (**Figure S1**).

### Source Reconstruction

Neural source activity was then estimated for evoked responses and average beta band activity using beamforming to achieve high spatial resolution. Each participant underwent structural MRI (Siemens Magnetom Prisma 3T, 1 mm isotropic MPRAGE). The pulse sequence parameters were as follows: 1 mm isotropic resolution; field of view, 256 × 240 mm; 192 slices; slice thickness, 1 mm; bandwidth per pixel, 290 Hz/pixel; flip angle, 9°; inversion time (TI), 1,100 ms; echo time (TE), 2.61 ms; and repetition time (TR), 2,300 ms. We furthermore acquired T2-weighted 3D images. The pulse sequence parameters were as follows: 1 mm isotropic resolution; field of view, 256 × 240 mm; 96 slices; slice thickness, 1.1 mm; bandwidth per pixel, 780 Hz/pixel; flip angle, 111°; echo time, 85 ms; and repetition time, 14,630 ms. These images were processed with FreeSurfer ^49,50^ to segment the head and brain surfaces. A volumetric source space (∼4,000 sources, 7.5 mm spacing) was generated using MNE-C’s watershed algorithm ^51^. Single compartment boundary element method (BEM) models were constructed, and MRI–MEG co-registration was performed using Polhemus FASTRAK fiducials and head shape points. Forward models were computed from co-registered data. A single-compartment BEM was used because MEG is relatively insensitive to conductivity differences between tissue compartments and is influenced primarily by primary neuronal currents rather than volume currents. Consequently, single-compartment models provide a robust and computationally efficient approximation for MEG source reconstruction ^52^.

Source estimation used a linearly constrained minimum variance (LCMV) beamformer ^53^ with the unit-noise-gain constraint, applied separately to evoked and beta-band data. The data covariance matrices were estimated from post-stimulus periods for the evoked responses and for the full epoch for the beta-band responses and used without regularization due to good conditioning of the covariance matrices. Magnetometer data were used for optimal sensitivity to deep sources.

### Statistical Analysis

The data met the assumptions of the statistical tests used in this study. Statistical analyses focused on identifying significant differences in beta-band responses between the ratios of the two omission conditions (non-jittered and jittered) in source space. Regions of interest were chosen based on previous experiments ^12,22^ and theory ^18^. The regions of interest were the left cerebellum crus I and left cerebellum VI, where we have found expectation related effects between non-jittered and jittered omissions in earlier studies ^12,22^ and the basal ganglia, here taken to be the caudate, the putamen and the pallidum (both hemispheres) ^18^. The selection of cerebellar and basal ganglia regions was hypothesis-driven and based on previous omission studies and theoretical accounts of temporal prediction ^12,22,39^. Within these predefined networks, source localization was data-driven, with statistical analyses focusing on the peak interaction effects identified within each region of interest.

Statistical maps for omissions were plotted for these regions, showcasing the interaction between Group (PD and controls) and Regularity (non-jittered and jittered), as these highlight where there are differences between the two groups in terms of how they build expectations under ideal (non-jittered) and less than ideal (jittered) conditions. The time range of interest was chosen to be from 100 ms before the expected stimulus to 100 ms *after* the expected stimulus. The −100 ms to 100 ms window was selected to capture neural dynamics immediately surrounding the expected but omitted stimulus, including both anticipatory (pre-omission) and immediate evaluative (post-omission) processes related to temporal prediction. This temporal window was additionally motivated by prior omission-related beta-band findings using the same paradigm ^12^. The difference between the non-jittered and the jittered conditions were calculated as the following ratio: 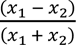. This measure expresses the difference between two conditions relative to their combined magnitude. Values close to zero indicate similar activity across conditions, whereas positive and negative values indicate relatively greater activity in one condition compared with the other.

For each of the two regions of interest (the cerebellum and the basal ganglia), we identified the source with the peak value for the interaction in the range from –100 ms to 100 ms. Although peak latencies and source locations are reported for descriptive purposes, statistical interpretation was restricted to effects supported by cluster-based permutation statistics, thereby reducing the likelihood that isolated local maxima reflected spurious findings. Confidence intervals on effect sizes were calculated using boot-strapping (5000 iterations).

To control the false alarm rate across the time dimension, we applied threshold-free cluster enhancement (TFCE) ^54^ to the time courses of the maximal sources in the cerebellum and basal ganglia. Because TFCE does not require specification of a fixed cluster-forming threshold, no initial cluster-defining threshold was applied. Clustering was performed across time points within each region of interest. TFCE was computed using a starting value of 0 and a step size of 0.1, with the standard parameter values E = 0.5 and H = 2. Statistical significance was estimated using 5,120 permutations.

To examine the relationship between neural activity and clinical severity, we computed partial correlations between beta-band activity and UPDRS-III scores within the PD group. Specifically, we calculated the ratio difference in beta power between non-jittered and jittered omissions at the peak latencies of –22 ms (cerebellum) and –15 ms (caudate) and tested their association with UPDRS-III motor scores. Age, LEDD, sex, and MDI were included as covariates to account for potential confounding effects, as these variables are known to influence neural oscillatory dynamics and motor performance in PD ^55–57^.

### Envelope correlations

To test functional connectivity between regions of interest, we used envelope correlation ^58^. We investigated the envelope correlations in the beta-band from –100 ms to 100 ms for the omitted stimulation. Regions of interest were based on the proposed network of sensory prediction from Andersen & Dalal (2024b) ^18^, which is constituted of basal ganglia, thalamus, cerebellum and the primary motor cortex. We included these areas bilaterally – and explored the functional connectivity within this proposed network. For descriptive purposes, we estimated the all-to-all connectivity between these nodes. Envelope correlation provides a non-directional measure of functional connectivity and does not permit inference regarding the direction of information flow between regions. To reduce the influence of signal leakage and zero-lag correlations, orthogonalized signals were used when estimating envelope correlations ^58,59^.

## 3. Results

As an initial validation of our data processing pipeline, we analyzed evoked responses in the primary and secondary somatosensory cortices (S1 & S2) across all participants (**Figure S1**). Clear somatosensory evoked responses were observed in both participants with PD and controls, with the expected SI (53 ms) and SII (132 ms) components following electrical stimulation.

These initial analyses were conducted separately for each group to assess whether the paradigm reproduced omission-related cerebellar beta-band effects previously reported in healthy participants before proceeding to group interaction analyses. In other words, we sought to replicate the findings of Andersen & Dalal (2021) ^12^ who reported increased beta-band activity (14-30 Hz) in Cerebellum-L VI for omissions following non-jittered compared with jittered somatosensory rhythms in neurotypical adult participants. Consistent with this earlier work, controls in the current study showed a significant effect in Cerebellum-L VI peaking at 85 ms: *t_24_* = 3.98, *p* = 0.00056, *d* = 0.80, (*d*_2.5%_ = 0.41; *d_97_*_.5%_ = 1.37), (MNI152: x = −15.0 mm, y = −75.0 mm, z = −23.0 mm) (**Figure 2**).

**Figure 2.**
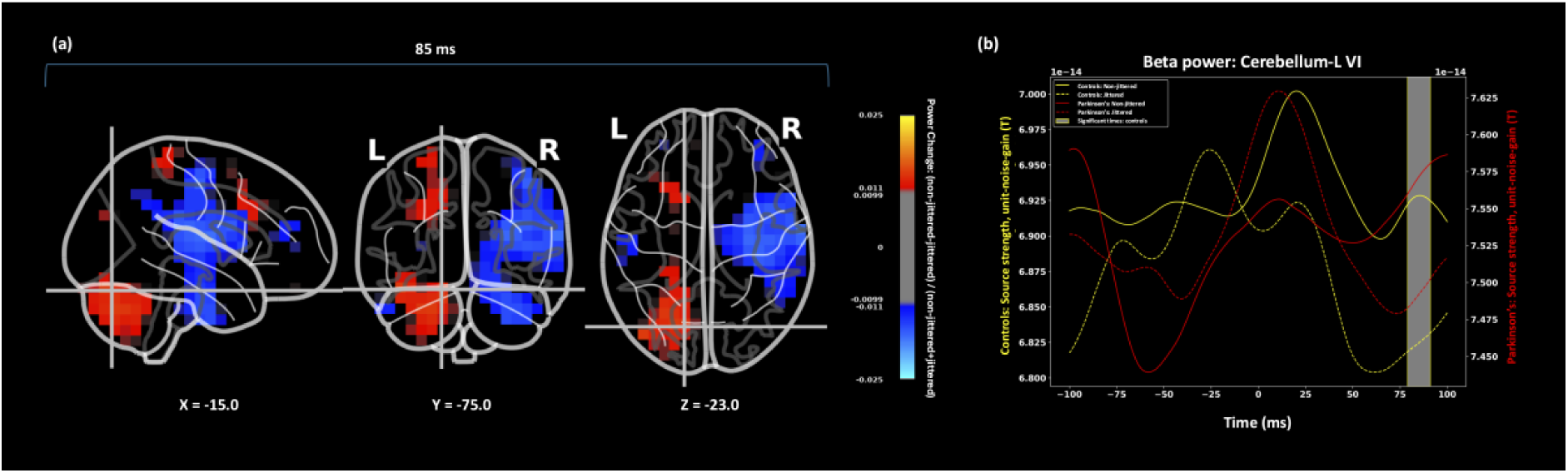
Replication of omission-related cerebellar beta-band effect in healthy controls. **(a)** Source-space map showing the effect of regularity (non-jittered vs. jittered) during omission processing in controls; **(b)** raw beta-band activity for controls and participants with PD during the jittered and non-jittered conditions. Gray box with yellow contour represents significant times for controls. Controls, n=25; Parkinson’s disease, n=24.

A cluster permutation test for the peak source in the Cerebellum-L VI for controls showed a significant difference between omissions following non-jittered and jittered rhythms (*p* = 0.047) The cluster informing the rejection of the null hypothesis extended from 79 ms to 91 ms. Andersen and Dalal (2021) ^12^ reported a corresponding cerebellar effect between 5 ms and 100 ms *after* the omission, falling within the previously reported interval, supporting replication of the earlier finding.

Applying the same analysis to participants with PD showed a peak difference at 49 ms: *t_23_* = 2.09, *p* = 0.048, *d* = 0.43 (*d*_2.5%_ = 0.019; *d_97_*_.5%_ = 0.97), (MNI152: x = −30.0 mm, y = ™38.0 mm, z = −38.0 mm). However, this effect did not survive cluster-based permutation correction (*p* = 0.438) and therefore should not be interpreted as evidence for a robust cerebellar regularity effect in the PD group.

When plotting the interaction (**Figure 3a**), a significant peak resulted in the time range from –100 ms to 100 ms in Cerebellum-L Crus I (cerebellar ROI) (MNI152: x = −52.5 mm, y = −67.5 mm, z = −37.5 mm; t = −22 ms), *F*_1,48_ = 12.0, *p* = 0.0011, *d* = −0.940, (*d*_2.5%_ = −1.54; *d_97_*_.5%_ = −0.43) (**Figure 3b**), and in Caudate-L (basal ganglia ROI) (MNI152: x = −15.0 mm, y = 15.0 mm, z = 0.0 mm; t = −15 ms), *F*_1,48_ = 19.5, *p* = 0.000056, *d* = 1.09, (*d*_2.5%_ = 0.57; *d_97_*_.5%_ = 1.74) (**Figure 3c**).

**Figure 3.**
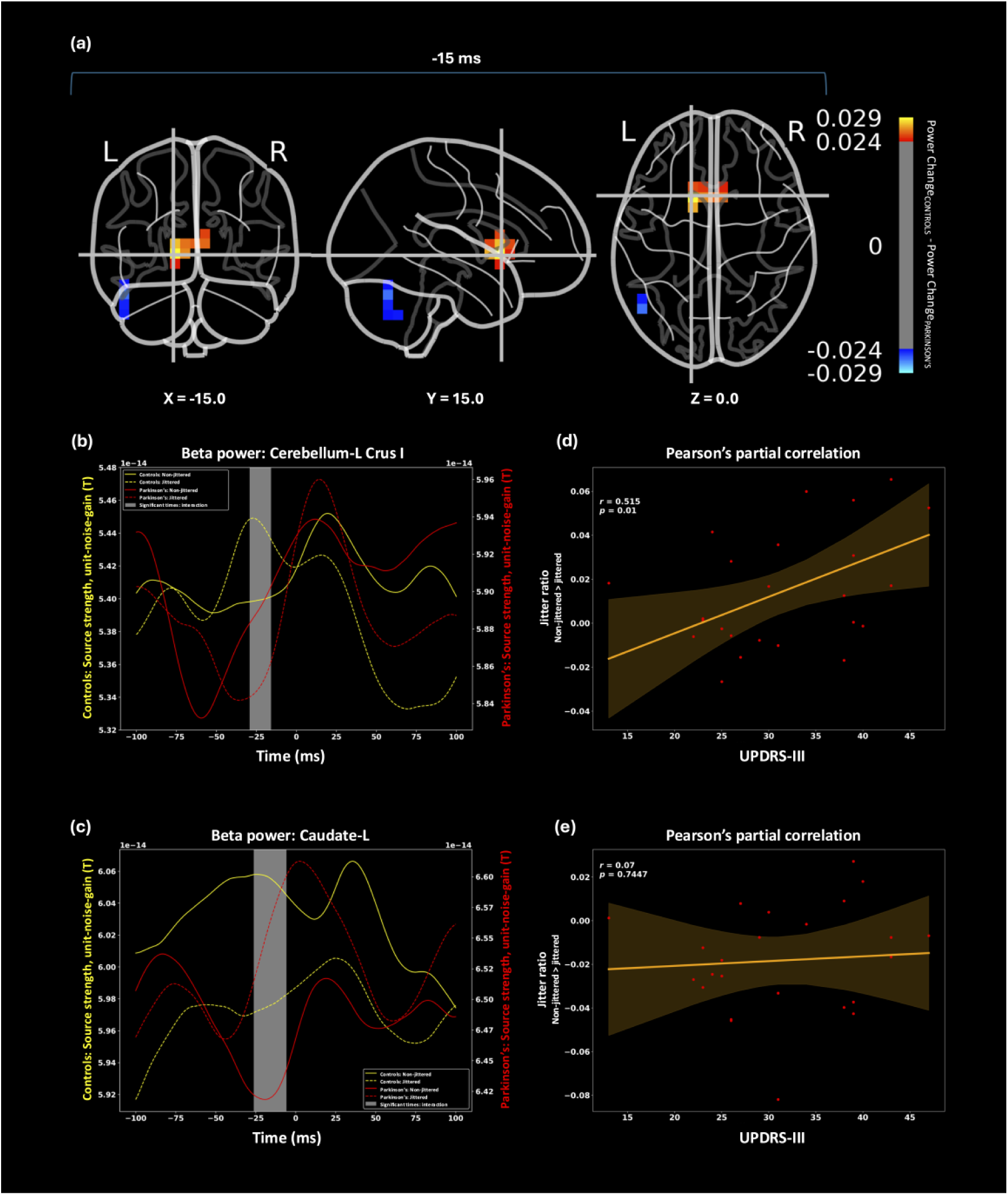
Interaction between Group (PD vs. controls) and Regularity (non-jittered vs. jittered). **(a)** Source-space map of the Group x Regularity interaction at the peak latency (t = –15 ms before the expected but omitted stimulus). Colors indicate differences in power changes (between non-jittered and jittered) between the groups (Controls minus PD). Prominent effects were observed in the left caudate and left cerebellum; **(b)** average beta-band activity in left cerebellum Crus I. Controls showed higher beta-band in the jittered condition than in the non-jittered condition, whereas participants with PD show the opposite pattern; **(c)** average beta-band activity in left caudate. Controls showed higher beta-band activity in non-jittered condition than in the jittered condition, whereas participants with PD showed the opposite pattern. Grey boxes indicate the temporal regions contributing to rejection of the cluster null hypothesis. Note the separate y-axes for controls and participants with PD; **(d)** partial correlation between UPDRS-III scores and the normalized beta-band contrast ((non-jittered − jittered)/(non-jittered + jittered)) extracted from the peak interaction source in left cerebellum Crus I at −22 ms; (**e**) partial correlation between UPDRS-III scores and the normalized beta-band contrast extracted from the peak interaction source in left caudate at –15 ms. Partial correlations were adjusted for age, sex, levodopa equivalent daily dose (LEDD), and Major Depression Inventory (MDI) scores. Shaded regions indicate 95% prediction intervals.Controls, n=25; Parkinson’s disease, n=24

For the left cerebellar Crus I, statistical inference identified significant time points from –29 ms to − 16 ms, peaking at –22 ms (TFCE-corrected, *p <* 0.05). A similar significant effect was observed for the left caudate, with significant time points spanning –26 ms to −6 ms, peaking at –15 ms (TFCE-corrected, *p <* 0.05) (see **Supplementary Information** for additional analyses).

When testing for the direction of the interaction for the Cerebellum-L Crus I (*t*=-22 ms), cerebellar beta-band source activity was higher for jittered than non-jittered omissions in controls, *t*_24_ = −2.30, *p* = 0.031, *d* = −0.46 (*d*_2.5%_ = −0.86; *d_97_*_.5%_ = −0.11), whereas the opposite pattern was found for participants with PD, *t*_23_ = 2.26, *p* = 0.034, *d* = 0.46 (*d*_2.5%_ = 0.081; *d_97_*_.5%_ = 0.88).

When testing for the direction of the interaction for Caudate-L (*t* = −15 ms), estimated caudate beta-band activity was higher for non-jittered than jittered omissions in controls, *t*_24_ = 3.51, *p* = 0.0018, *d* = 0.70 (*d*_2.5%_ = 0.44; *d_97_*_.5%_ = 1.04), whereas the opposite pattern was found for participants with PD, *t*_23_ = −2.08, *p* = 0.048, *d* = 0.67 (*d*_2.5%_ = −0.91; *d_97_*_.5%_ = −0.050).

To test the link between PD related symptoms and the reported interaction in the cerebellum (*r* = 0.452, *p* = 0.0267) (**Figure 3d**) and the basal ganglia (*r* = 0.287, *p* = 0.174) (**Figure 3e**), we calculated the partial correlation between the peak ratio difference, at –22 ms and –15 ms respectively, for each PD participant with the UPDRS-III score, after regressing out age, sex, MDI, and LEDD.

In summary, controls showed differential cerebellar beta-band activity in response to omissions following non-jittered versus jittered, peaking at 85 ms *after* the expected but omitted stimulus, replicating earlier findings ^12^. In contrast, *before* the expected but omitted stimulus, beta band activity in the Cerebellum-L Crus I (−22 ms) and the basal ganglia (Caudate-L) (−15 ms) differed as a function of both group (PD vs controls) and rhythmic regularity (non-jittered vs jittered), suggesting altered temporal prediction-related processing in PD. These interaction effects were estimated using the ratio between non-jittered and jittered omissions (**Figure 3d** and **Figure 3e**).

To explore, whether the envelope analysis supported an interaction between Group and Regularity, we chose Caudate-L as a seed and included bilaterally, the cerebellar regions found in the current analysis (Cerebellum-L VI and Cerebellar-L Crus I) and the regions specified in earlier work (Motor Cortex and Thalamus and Caudate) ^18^. This was, in part, motivated by the descriptive plot (**Figure 4a**) showing that the greatest differences were connections involving the caudate, and, in part, motivated by the *a priori* knowledge that the basal ganglia should be involved due to PD. By running an ANOVA testing for the interaction between Group and Regularity while controlling for Region of Interest (10 levels), we found the following significant interaction: *F_1,967_* = 4.79, *p* = 0.0289 **(Figure 4b**) Specifically, participants with PD showed greater connectivity during the jittered condition relative to the non-jittered condition, whereas controls showed a smaller difference between conditions. Again, these connectivity analyses were exploratory and intended to characterize network-level differences associated with rhythmic regularity.

**Figure 4.**
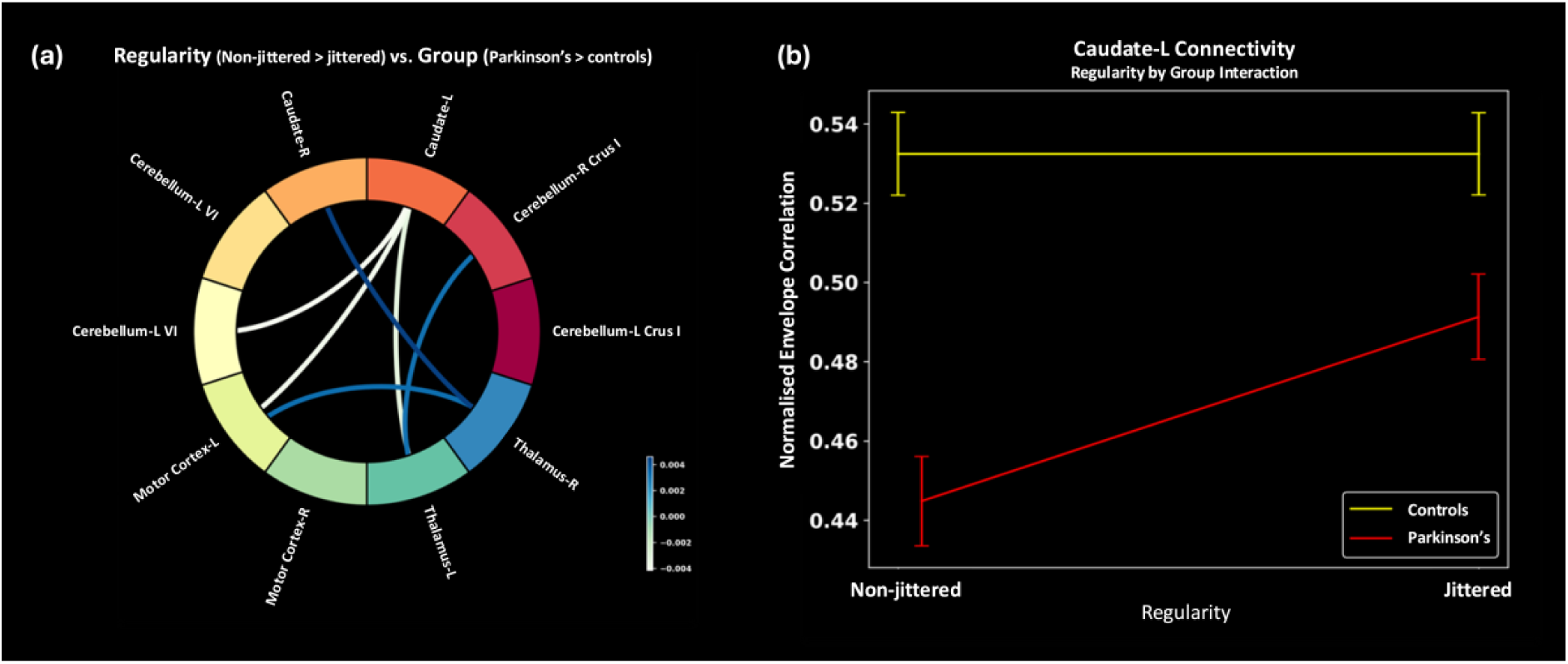
Functional Connectivity. Envelope correlation findings: **(a)** all-to-all differences in connectivity between regions of interest. Only the six largest interaction effects are displayed for visualization purposes, representing the minimum number of connections required to show all functional links involving the left caudate within the proposed cerebello-striatal-thalamo- cortical network (Andersen & Dalal, 2024b). The left caudate showed connectivity differences with all regions of the proposed network, i.e. cerebellum, thalamus and the motor cortex. **(b)** Group by Regularity interaction for the connections involving the left caudate. Participants with PD show higher connectivity during the jittered condition than during the non-jittered condition. Error bars are standard errors of the mean. Controls, n=25; Parkinson’s disease, n=24.

## 4. Discussion

### Summary of Key Findings

This study investigated cerebellar and basal ganglia beta-band (14-30 Hz) activity in participants with PD compared to controls during a rhythmic somatosensory timing task. Our findings show that:

1. *After* the expected but omitted stimulus (85 ms), controls exhibited a cerebellar beta-band regularity effect, with larger responses to omissions following regular (non-jittered) than irregular (jittered) somatosensory rhythms, replicating previous evidence for cerebellar involvement in temporal evaluation ^12^.
2. *Before* the expected but omitted stimulus, beta-band activity in both the cerebellum (−22 ms) and the basal ganglia (−15 ms) showed significant group-by-regularity interactions. Controls and participants with PD exhibited opposing regularity effects, with PD participants showing relatively greater responses to irregular than regular rhythms, consistent with altered temporal prediction-related processing.
3. In participants with PD, larger cerebellar beta-band differences between regular and irregular rhythms *before* the expected but omitted stimulus (−22 ms) were associated with higher motor symptom severity (UPDRS-III), demonstrating an association between altered temporal prediction-related activity and motor symptom severity.
4. Participants with PD exhibited greater caudate-centered beta-band connectivity during irregular than regular rhythms, which may reflect altered engagement of the cerebello–basal ganglia–thalamo–cortical network when temporal regularity is reduced.

### Explication of interactions

To clarify the group interactions, we first consider the control group. Consistent with theories of rhythmic cognition, basal ganglia activity differentiated between regular and irregular rhythms, exhibiting higher beta-band activity for non-jittered than jittered rhythms, consistent with previous functional magnetic resonance imaging (fMRI) findings ^11,60^.

In terms of the cerebellar activity *before* the expected stimulus, beta-band power was higher following jittered than non-jittered rhythms. One possible explanation is that the cerebellum shows reduced activity for non-jittered rhythms because the timing of the upcoming, expected, but omitted stimulus is highly predictable. This interpretation aligns with findings suggesting that the cerebellum is more engaged when temporal regularity is reduced or timing prediction is less reliable, whereas the basal ganglia are more engaged by temporally regular input ^11,17^.

In PD, this pattern was reversed. Basal ganglia activity was higher for jittered than non-jittered rhythms, possibly reflecting increased reliance on internally generated temporal predictions and network engagement to extract temporal structure when external timing cues are less reliable ^11^. Conversely, cerebellar activity in PD was higher for non-jittered than jittered rhythms, suggesting altered cerebellar recruitment: when basal ganglia processing of temporal regularity is altered, the cerebellum may assume a greater role in supporting temporal predictions.

An alternative interpretation is that the observed group differences do not solely reflect a redistribution of processing between basal ganglia and cerebellar systems. Dopaminergic medication ^61^ and broader alterations in oscillatory dynamics associated with PD ^62^, may also contribute to the observed effects. In addition, non-specific factors such as attention and vigilance cannot be completely excluded ^63^, although participants were instructed to watch a nature documentary and no behavioral task was required. Consequently, the present findings should be interpreted as being consistent with altered cerebello–basal ganglia contributions to temporal prediction rather than as definitive evidence of compensatory cerebellar recruitment.

Although these pre-omission effects occurred immediately *before* the expected but omitted stimulus and were sensitive to temporal regularity, they should not be interpreted as direct measures of temporal prediction. Given the relatively long inter-stimulus interval (1,497 ms), the observed beta-band activity may also reflect anticipatory neural dynamics or residual processing associated with the preceding stimulus sequence ^63^. The present interpretation is therefore based on the temporal proximity of these effects to the expected stimulus and their modulation by temporal regularity, but alternative explanations cannot be excluded.

### Cerebellum and temporal processing: *Evaluation* vs. *Prediction*

Our findings show that the cerebellum displays a beta-band response peaking at 85 ms *after* the omitted stimulus in controls. This replicates earlier findings ^12^ and is consistent with the proposal by Andersen & Dalal ^18^ that the cerebellum plays a role in integrating sensory predictions with sensory feedback. The present findings are also consistent with previous evidence that the cerebellum exhibits beta-band activity during sensorimotor and predictive processing. Cerebellar beta oscillations have been reported in MEG and EEG studies of somatosensory prediction and omission processing, as well as in studies of motor control and temporal processing, supporting the physiological plausibility of cerebellar beta-band involvement in the present task ^12,16,18,22,31,32,39^.

Although PD pathophysiology is often associated particularly with alterations in lower beta frequencies ^64^, the broader 14–30 Hz range examined here is consistent with our previous studies of cerebellar temporal prediction and omission processing, and there was no *a priori* basis for restricting cerebellar effects to the lower beta range ^18,22,39,65^.

The reconstruction of the post-omission and pre-omission effects to different cerebellar subregions may further reflect functional specialization within the cerebellum ^66^. Cerebellum VI has frequently been associated with sensorimotor and somatosensory processing, consistent with the evaluative responses observed *after* the omitted stimulus. In contrast, cerebellum Crus I has been linked to higher-order predictive and integrative functions, including the coordination of temporally structured information across distributed networks ^67,68^. The pre-omission interaction effects observed in Crus I may therefore reflect anticipatory predictive processing related to upcoming sensory events rather than sensory evaluation following omission ^16^.

However, the observed differences in cerebellum Crus I *before* the omitted stimulus suggest altered cerebellar involvement in temporal prediction-related processing in PD. This interpretation is supported by the association between symptom severity and the magnitude of the cerebellar responses to non-jittered versus jittered omissions.

Our findings contribute to a growing body of literature suggesting that the cerebellum plays a role in building sensory predictions through internal forward models ^69–71^, although the extent to which cerebellar computations reflect temporal prediction, timing mechanisms, or an interaction between the two remains debated ^71,72^. The observed differences in cerebellar regions in participants with PD *before* the omitted stimulus may reflect distinct recruitment of cerebellar predictive mechanisms in response to altered basal ganglia function, consistent with previous interpretations of increased cerebellar engagement in PD as a potential compensatory or adaptive network response ^72,73^. Importantly, greater cerebellar alterations were associated with higher UPDRS-III scores, suggesting that these neural differences are related to clinical symptom severity. Together, these findings support the idea that cerebellar involvement in PD reflects altered network engagement during temporal prediction, although the present data do not establish whether such recruitment is functionally beneficial or compensatory ^74^.

It should be noted that the seventh event was never globally expected, as each stimulation train consisted of six pulses. However, prior work has shown that omissions can generate cerebellar responses even when they are not globally expected, provided they are locally predictable within a train ^12,16,22^. Our paradigm thus probes local temporal predictions rather than global sequence expectations, consistent with the replicated cerebellar omission responses reported here.

### Basal Ganglia Dysfunction and Altered Timing Mechanisms in Parkinson’s disease

The significant alteration in beta-band activity in the caudate nucleus between participants with PD and controls underscores the basal ganglia’s pivotal role in temporal processing. The basal ganglia, particularly the caudate and putamen, have long been implicated in the regulation of motor control and timing ^9^. Our findings are consistent with a large body of literature showing that PD impairs both motor and perceptual timing tasks. Notably, the basal ganglia dysfunction plays a central role in these deficits ^30^.

Beta-band activity within the basal ganglia is thought to maintain the current motor state and suppress voluntary movements until the appropriate time for action ^75^. This rhythmic activity is crucial for the fine-tuning of temporal predictions and the execution of time-sensitive motor behaviors ^61^. The different beta-band responses observed in PD suggest a failure in the basal ganglia’s ability to appropriately modulate these timing signals.

Furthermore, neuroimaging studies have shown that the basal ganglia are critical for interval timing and the perception of temporal sequences ^29,76,77^. These studies have shown that the caudate nucleus is particularly involved in the processing of temporal expectations and the updating of internal models based on sensory feedback. Our findings extend this understanding by showing that participants with PD exhibit altered beta-band activity in the caudate during tasks requiring precise temporal predictions, suggesting altered temporal prediction-related processing within basal ganglia networks in PD.

The interpretation of caudate-related findings in PD should nevertheless be considered cautiously. In early PD, dopaminergic depletion is often more pronounced in dorsal striatal regions such as the putamen than in more ventral striatal territories ^78,79^. Because participants in the present study varied in disease duration and were tested in the ON-medication state, some of the observed caudate-related differences may additionally reflect differential dopaminergic modulation of relatively preserved striatal circuits ^80,81^.

### Beta-Band Connectivity between Basal Ganglia and Cerebellum, Thalamus and Motor Cortex

We find evidence that the connectivity in the beta-band between the caudate and cerebellum, thalamus and motor cortex specifically changes in PD. Compared to controls, participants with PD showed an increase in connectivity for the jittered condition compared to the non-jittered condition. Similarly, estimated caudate source activity was higher for jittered than non-jittered stimuli in PD. Importantly, the connectivity analysis identifies connections exhibiting the strongest differences between groups rather than providing a complete description of the underlying functional network. Therefore, the prominence of connections centered in the caudate suggests that PD alterations are most pronounced in pathways involving the basal ganglia. The absence of equally strong differences between other nodes should not be interpreted as evidence that these regions are not functionally connected, but rather that those connections contributed less to the observed group effect. Within this context, increased coupling may reflect tighter coordination between basal ganglia and connected cerebellar, thalamic, and cortical regions integrating timing information with action plans (Andersen & Dalal, 2024a). In other words, reduced temporal regularity in the jittered condition may increase the computational demands placed on the cerebello–striatal–thalamo–cortical network, requiring stronger communication to maintain internal temporal predictions despite less reliable external timing cues (Grahn & Rowe, 2009). Such increased coupling may reflect a greater reliance on internally generated timing mechanisms when external rhythmic structure becomes less predictable ^17,82–84^.

Importantly, these connectivity findings should be interpreted cautiously. Envelope correlation provides a non-directional estimate of functional connectivity and cannot determine the direction of information flow between regions. Moreover, although orthogonalization reduces the influence of signal leakage and zero-lag correlations ^58,59^, the observed effects may also reflect broader changes in beta-band dynamics rather than specific communication between individual nodes of the network.

Our connectivity findings also align with extensive work on motor cortical beta oscillations. Motor cortex beta activity is thought to maintain the current motor set and inhibit premature actions, with transient beta suppression enabling movement initiation ^61,75^. In PD, exaggerated beta synchrony between motor cortex and basal ganglia is a well-documented pathophysiological hallmark ^85^ and is attenuated by dopaminergic therapy or deep brain stimulation. In our study, participants with PD showed an increase in connectivity during the jittered condition relative to the non-jittered condition. We interpret this increase as potentially reflecting the same pathophysiological mechanism of excessive beta synchrony, but here manifesting in a context where the network relies on additional resources to impose regularity on irregular input. Thus, the altered connectivity we observe between caudate, cerebellum, thalamus and motor cortex likely reflects disruptions in this wider beta-band network, consistent with both cortical and subcortical contributions to impaired temporal prediction and motor control in PD. The seeming difference in connectivity between the two groups may be driven by controls in general showing higher beta band activity than PD participants (**Figure 3d** and **Figure3e**) ^86^. However, this is not an issue for the interpretation of the interaction, which is independent of average differences between groups.

Taken together, the present findings suggest that temporal *prediction* and temporal *evaluation* involve partially overlapping but distinct contributions from cerebellar and basal ganglia systems. The post-omission cerebellar response observed in controls is consistent with a role in evaluating predicted sensory events, whereas the pre-omission effects in both cerebellum and basal ganglia indicate involvement in generating or maintaining temporal expectations before an expected event occurs. The altered connectivity observed in PD further suggests that these functions depend on coordinated interactions across a broader cerebello–striatal–thalamo–cortical network rather than on isolated computations within individual structures.

### Limitations and future perspectives

One limitation concerns the participant sample. Although the sample size is comparable to several task-based MEG studies in Parkinson’s disease, the study may still be underpowered to detect smaller effects, particularly for exploratory connectivity analyses and subgroup-related variability.

Also, our study included only medicated participants with PD, meaning that the effects of dopaminergic medication on beta-band activity cannot be excluded. Prior work has shown that levodopa may reduce beta-band synchrony within basal ganglia–cortical networks ^87^, although the extent to which these effects apply specifically to caudate-related activity remains unclear. Consequently, some of the observed group differences may reflect a combination of disease-related and medication-related influences on beta-band dynamics. To partially address this issue, we additionally controlled for LEDD in the correlation analyses, although future ON/OFF medication designs will be necessary to disentangle medication effects from disease-related neural alterations more directly. Additionally, disease duration varied across participants with PD, and the study was not designed to stratify participants according to disease stage or medication-response profiles ^81^.

Another limitation is that despite MEG being a powerful tool for investigating neural activity with high temporal resolution, it has limited sensitivity to central brain structures. MEG sensitivity to deep structures such as the basal ganglia remains limited, and source localization should therefore be interpreted cautiously. This limitation is mitigated by the observation that our magnetoencephalographic basal ganglia findings in controls concord with what is found in functional magnetic resonance imaging studies ^11,17,77^, namely that the basal ganglia preferentially respond to non-jittered stimuli. It is furthermore mitigated by the *a priori* knowledge that this is exactly where participants with PD should show distinct activity, which is precisely what our results present. Notably, as we relied on omissions to elicit activity, cortical activity of the cerebrum, which might otherwise mask sub-cortical activity, was kept to a minimum ^33^. Although converging computational and empirical evidence suggests that subcortical contributions may be detectable with MEG under specific conditions, source localization in deep brain structures should nevertheless be interpreted cautiously. Accordingly, the present findings should be understood as source-reconstructed estimates consistent with basal ganglia involvement rather than direct recordings from these structures.

Although participants with PD exhibited distinct patterns of beta-band activity and connectivity, the present findings do not, by themselves, establish how these neural alterations relate to temporal prediction or motor performance. An important next step will be to combine omission paradigms with behavioral measures of temporal prediction and motor timing, as recently demonstrated by Andersen et al. (2024) ^22^, allowing neural signatures to be directly linked to behavioral performance. Such studies will help determine the functional significance of the observed neural alterations and clarify how they contribute to predictive processing in PD.

An important consideration concerns the lateralization of our findings. The most robust source-level effects emerged in the left cerebellum and left basal ganglia, despite the well-documented contralateral connectivity of the cerebello–cortico–striatal system (i.e., left cerebellum to right basal ganglia and motor cortex). Several factors may contribute: (i) tactile stimulation was delivered to the right index finger, favoring contralateral (left-hemisphere) responses; (ii) all participants were right-handed, consistent with left-dominant sensorimotor processing; and (iii) MEG beamforming sensitivity can differ across hemispheres. Importantly, functional asymmetries in cerebellar and basal ganglia activation during timing tasks have been reported previously ^10,25,88,89^, suggesting that lateralized effects are not unusual. At the connectivity level, we additionally observed coupling between the left basal ganglia and right cerebellum (**Figure 4**), consistent with cross-hemispheric cerebello–striatal loops. Taken together, the apparent “left–left” effects may therefore reflect both methodological sensitivity and known asymmetries, but our sample was not stratified by side of symptom onset in PD, so we cannot directly relate lateralization to clinical laterality. Future work with larger and side-stratified samples may clarify this issue. No specific hypotheses were made regarding individual basal ganglia subregions or hemispheric lateralization, and such findings should therefore be interpreted cautiously.

It should be noted that there were differences in sex, age and MDI. To mitigate the effects of these differences, we conducted complementary analyses controlling for these. This importantly did not substantially alter the reported interaction effects.

### Conclusion

In conclusion, participants with PD showed evaluative cerebellar beta-band responses alongside altered predictive timing-related signals involving the basal ganglia. Increased cerebellar activity *before* predicted stimuli was associated with symptom severity and is consistent with altered cerebello-basal ganglia contributions to temporal prediction. PD also disrupted connectivity within a sensory-integration network involving the cerebellum, basal ganglia, thalamus and primary motor cortex.

These findings extend beyond clinical characterization of PD: they provide mechanistic insight into how rhythmic cognition emerges from the interplay between predictive and evaluative processes in subcortical–cortical networks. The present findings suggest that beta-band dynamics within the cerebello–striatal–thalamo–cortical circuit may support temporal prediction during regular rhythms and evaluation when these predictions are violated. In PD, basal ganglia dysfunction seems to shift this balance, increasing reliance on cerebellar engagement and altering network connectivity, particularly under irregular rhythmic conditions.

By combining high-temporal-resolution MEG with a rhythmic omission paradigm, we identify key neural signatures of predictive timing and their reorganization in PD. This work bridges basic rhythmic cognition research and clinical neuroscience, offering a network-level framework to interpret timing dysfunction in PD.

## Author Contributions

VPN: Investigation, Writing – Original Draft, Writing – Review & Editing, Visualization.

LMA: Conceptualization, Methodology, Software, Validation, Formal Analysis, Investigation, Data Curation, Writing – Original Draft, Writing – Review & Editing, Visualization, Supervision, Project Administration, Funding Acquisition.

## Acknowledgements

V.P.N. is funded by the Center for Music in the Brain (MIB), supported by the Lundbeck Foundation (R469-2024-1573) and Købmand Herman Sallings Fond. L.M.A. was funded by the Lundbeck Foundation (R322-2019-1841). The funders had no role in study design, data collection and analysis, decision to publish, or preparation of the manuscript. The authors declare that there are no conflicts of interest relevant to this work, and that there are no additional disclosures to report.

## Data Availability

The raw MRI data underlying this study contain sensitive personal information and cannot be fully shared publicly due to restrictions imposed by the General Data Protection Regulation (GDPR) and Danish national data protection laws. The data and materials that can be shared publicly are available at https://github.com/ualsbombe/pd_cerebellum. Requests for access to the raw MRI data can be submitted to the corresponding author and require a formal data-sharing agreement and approval from the relevant parties. The raw MEG data are available upon reasonable request to the corresponding author.

## Code Availability

The analysis scripts and code used for data processing, statistical analysis, and visualization are publicly available at https://github.com/ualsbombe/pd_cerebellum.

## References

1. Blakemore, S.-J., Wolpert, D. M. & Frith, C. D. Central cancellation of self-produced tickle sensation. Nat. Neurosci. 1, 635–640 (1998).

2. Clark, A. Whatever next? Predictive brains, situated agents, and the future of cognitive science. Behav. Brain Sci. 36, 181–204 (2013).

3. Gross, J. et al. The neural basis of intermittent motor control in humans. Proc. Natl. Acad. Sci. 99, 2299–2302 (2002).

4. Miall, R. C., Weir, D. J., Wolpert, D. M. & Stein, J. F. Is the cerebellum a smith predictor? J. Mot. Behav. 25, 203–216 (1993).

5. Wolpert, D. M. & Kawato, M. Multiple paired forward and inverse models for motor control. Neural Netw. Off. J. Int. Neural Netw. Soc. 11, 1317–1329 (1998).

6. Schmahmann, J. D. & Pandya, D. N. The cerebrocerebellar system. Int. Rev. Neurobiol. 41, 31–60 (1997).

7. Kakei, S. et al. Contribution of the Cerebellum to Predictive Motor Control and Its Evaluation in Ataxic Patients. Front. Hum. Neurosci. 13, (2019).

8. Peterburs, J. & Desmond, J. E. The role of the human cerebellum in performance monitoring. Curr. Opin. Neurobiol. 40, 38–44 (2016).

9. Coull, J. T., Hwang, H. J., Leyton, M. & Dagher, A. Dopamine Precursor Depletion Impairs Timing in Healthy Volunteers by Attenuating Activity in Putamen and Supplementary Motor Area. J. Neurosci. 32, 16704–16715 (2012).

10. Grahn, J. A. & Brett, M. Rhythm and beat perception in motor areas of the brain. J. Cogn. Neurosci. 19, 893–906 (2007).

11. Grahn, J. A. & Rowe, J. B. Feeling the Beat: Premotor and Striatal Interactions in Musicians and Nonmusicians during Beat Perception. J. Neurosci. 29, 7540–7548 (2009).

12. Andersen, L. M. & Dalal, S. S. The cerebellar clock: Predicting and timing somatosensory touch. NeuroImage 238, 118202 (2021).

13. Brochard, R., Touzalin, P., Després, O. & Dufour, A. Evidence of beat perception via purely tactile stimulation. Brain Res. 1223, 59–64 (2008).

14. Buhusi, C. V. & Meck, W. H. What makes us tick? Functional and neural mechanisms of interval timing. Nat. Rev. Neurosci. 6, 755–765 (2005).

15. Grube, M., Cooper, F. E., Chinnery, P. F. & Griffiths, T. D. Dissociation of duration-based and beat-based auditory timing in cerebellar degeneration. Proc. Natl. Acad. Sci. U. S. A. 107, 11597–11601 (2010).

16. Tesche, C. D. & Karhu, J. J. T. Anticipatory cerebellar responses during somatosensory omission in man. Hum. Brain Mapp. 9, 119–142 (2000).

17. Teki, S., Grube, M., Kumar, S. & Griffiths, T. D. Distinct Neural Substrates of Duration-Based and Beat-Based Auditory Timing. J. Neurosci. 31, 3805–3812 (2011).

18. Andersen, L. M. & Dalal, S. S. The role of the cerebellum in timing. Curr. Opin. Behav. Sci. 59, 101427 (2024).

19. Criscuolo, A., Schwartze, M., Nozaradan, S. & Kotz, S. A. Basal ganglia and cerebellar lesions causally impact the neural encoding of temporal regularities. Imaging Neurosci. 3, imag_a_00492 (2025).

20. Caligiore, D. et al. Parkinson’s disease as a system-level disorder. NPJ Park. Dis. 2, 16025 (2016).

21. Grahn, J. A. & Brett, M. Impairment of beat-based rhythm discrimination in Parkinson’s disease. Cortex 45, 54–61 (2009).

22. Andersen, L. M. & Dalal, S. S. Detection of Threshold-Level Stimuli Modulated by Temporal Predictions of the Cerebellum. eneuro 11, ENEURO.0070-24.2024 (2024).

23. Arnal, L. H. Predicting “When” Using the Motor System’s Beta-Band Oscillations. Front. Hum. Neurosci. 6, (2012).

24. te Woerd, E. S., Oostenveld, R., Bloem, B. R., de Lange, F. P. & Praamstra, P. Effects of rhythmic stimulus presentation on oscillatory brain activity: the physiology of cueing in Parkinson’s disease. NeuroImage Clin. 9, 300–309 (2015).

25. Vinding, M. C. et al. Attenuated beta rebound to proprioceptive afferent feedback in Parkinson’s disease. Sci. Rep. 9, 2604 (2019).

26. Vinding, M. C. et al. Oscillatory and non-oscillatory features of the magnetoencephalic sensorimotor rhythm in Parkinson’s disease. Npj Park. Dis. 10, 1–13 (2024).

27. DeLong, M. R. & Wichmann, T. Circuits and circuit disorders of the basal ganglia. Arch. Neurol. 64, 20–24 (2007).

28. Kalia, L. V. & Lang, A. E. Parkinson’s disease. Lancet Lond. Engl. 386, 896–912 (2015).

29. Jones, C. R. G. & Jahanshahi, M. Motor and Perceptual Timing in Parkinson’s Disease. in Neurobiology of Interval Timing (eds Merchant, H. & de Lafuente, V.) 265–290 (Springer, New York, NY, 2014). doi:10.1007/978-1-4939-1782-2_14.

30. Harrington, D. L. et al. Neurobehavioral Mechanisms of Temporal Processing Deficits in Parkinson’s Disease. PLOS ONE 6, e17461 (2011).

31. Andersen, L. M., Jerbi, K. & Dalal, S. S. Can EEG and MEG detect signals from the human cerebellum? NeuroImage 215, 116817 (2020).

32. Samuelsson, J. G., Sundaram, P., Khan, S., Sereno, M. I. & Hämäläinen, M. S. Detectability of cerebellar activity with magnetoencephalography and electroencephalography. Hum. Brain Mapp. 41, 2357–2372 (2020).

33. Krishnaswamy, P., aet al. Sparsity enables estimation of both subcortical and cortical activity from MEG and EEG. Proc. Natl. Acad. Sci. 114, E10465–E10474 (2017).

34. Attal, Y., Maess, B., Friederici, A. & David, O. Head models and dynamic causal modeling of subcortical activity using magnetoencephalographic/electroencephalographic data. 23, 85–95 (2012).

35. Fujioka, T., Trainor, L. J., Large, E. W. & Ross, B. Internalized Timing of Isochronous Sounds Is Represented in Neuromagnetic Beta Oscillations. J. Neurosci. 32, 1791–1802 (2012).

36. Piastra, M. C. et al. A comprehensive study on electroencephalography and magnetoencephalography sensitivity to cortical and subcortical sources. Hum. Brain Mapp. 42, 978–992 (2021).

37. Li, J., Zhai, M., Liu, L., Qian, Y. & Liao, Y. Beta Oscillations as Internal Timers: Independent Contributions of Temporal Predictability to Motor Preparation. Psychophysiology 63, e70249 (2026).

38. Postuma, R. B. et al. MDS clinical diagnostic criteria for Parkinson’s disease. Mov. Disord. 30, 1591–1601 (2015).

39. Andersen, L. M. & Lundqvist, D. Somatosensory responses to nothing: An MEG study of expectations during omission of tactile stimulations. NeuroImage 184, 78–89 (2019).

40. McCusker, M. C. et al. Altered neural oscillations during complex sequential movements in patients with Parkinson’s disease. NeuroImage Clin. 32, 102892 (2021).

41. Vinding, M. C. et al. Reduction of spontaneous cortical beta bursts in Parkinson’s disease is linked to symptom severity. Brain Commun. 2, fcaa052 (2020).

42. Nasreddine, Z. S., et al. The Montreal Cognitive Assessment, MoCA: a brief screening tool for mild cognitive impairment. J Am Geriatr Soc 53, 695–699 (2005).

43. Olsen, L. R., Jensen, D. V., Noerholm, V., Martiny, K. & Bech, P. The internal and external validity of the Major Depression Inventory in measuring severity of depressive states. https://doi.org/10.1017/S0033291702006724 (2003) doi:10.1017/S0033291702006724.

44. Goetz, C. G. et al. Movement Disorder Society-sponsored revision of the Unified Parkinson’s Disease Rating Scale (MDS-UPDRS): Scale presentation and clinimetric testing results. Mov. Disord. 23, 2129–2170 (2008).

45. Tomlinson, C. L. et al. Systematic review of levodopa dose equivalency reporting in Parkinson’s disease. Mov. Disord. 25, 2649–2653 (2010).

46. Peirce, J. W. Generating stimuli for neuroscience using PsychoPy. *Front*. Neuroinformatics 2, (2009).

47. Gramfort, A. et al. MEG and EEG data analysis with MNE-Python. Front. Neurosci. 7, (2013).

48. Andersen, L. M. Group Analysis in MNE-Python of Evoked Responses from a Tactile Stimulation Paradigm: A Pipeline for Reproducibility at Every Step of Processing, Going from Individual Sensor Space Representations to an across-Group Source Space Representation. Front. Neurosci. 12, (2018).

49. Dale, A. M., Fischl, B. & Sereno, M. I. Cortical surface-based analysis. I. Segmentation and surface reconstruction. NeuroImage 9, 179–194 (1999).

50. Fischl, B. FreeSurfer. NeuroImage 62, 774–781 (2012).

51. Gramfort, A. et al. MNE software for processing MEG and EEG data. NeuroImage 86, 446–460 (2014).

52. Hämäläinen, M., Hari, R., Ilmoniemi, R. J., Knuutila, J. & Lounasmaa, O. V. Magnetoencephalography---theory, instrumentation, and applications to noninvasive studies of the working human brain. Rev. Mod. Phys. 65, 413–497 (1993).

53. Sekihara, K., Hild, K. E., Dalal, S. S. & Nagarajan, S. S. Performance of prewhitening beamforming in MEG dual experimental conditions. IEEE Trans. Biomed. Eng. 55, 1112–1121 (2008).

54. Maris, E. & Oostenveld, R. Nonparametric statistical testing of EEG- and MEG-data. J. Neurosci. Methods 164, 177–190 (2007).

55. Abdullah, R. et al. Parkinson’s disease and age: The obvious but largely unexplored link. Exp. Gerontol. 68, 33–38 (2015).

56. Levy, G. et al. Contribution of Aging to the Severity of Different Motor Signs in Parkinson Disease. Arch. Neurol. 62, 467–472 (2005).

57. Seidler, R. D. et al. Motor control and aging: Links to age-related brain structural, functional, and biochemical effects. Neurosci. Biobehav. Rev. 34, 721–733 (2010).

58. Hipp, J. F., Hawellek, D. J., Corbetta, M., Siegel, M. & Engel, A. K. Large-scale cortical correlation structure of spontaneous oscillatory activity. Nat. Neurosci. 15, 884–890 (2012).

59. Brookes, M. J., Woolrich, M. W. & Barnes, G. R. Measuring functional connectivity in MEG: A multivariate approach insensitive to linear source leakage. NeuroImage 63, 910–920 (2012).

60. Grahn, J. A. & Rowe, J. B. Finding and feeling the musical beat: Striatal dissociations between detection and prediction of regularity. Cereb. Cortex 23, 913–921 (2013).

61. Jenkinson, N. & Brown, P. New insights into the relationship between dopamine, beta oscillations and motor function. Trends Neurosci. 34, 611–618 (2011).

62. Hammond, C., Bergman, H. & Brown, P. Pathological synchronization in Parkinson’s disease: networks, models and treatments. Trends Neurosci. 30, 357–364 (2007).

63. Kok, P., Rahnev, D., Jehee, J. F. M., Lau, H. C. & de Lange, F. P. Attention Reverses the Effect of Prediction in Silencing Sensory Signals. Cereb. Cortex 22, 2197–2206 (2012).

64. Yu, Y. et al. Parkinsonism Alters Beta Burst Dynamics across the Basal Ganglia–Motor Cortical Network. J. Neurosci. 41, 2274–2286 (2021).

65. Hari, R. & Puce, A. MEG - EEG Primer. (Oxford University Press, 2023). doi:10.1093/med/9780197542187.001.0001.

66. Stoodley, C. J. & Schmahmann, J. D. Functional topography in the human cerebellum: a meta-analysis of neuroimaging studies. NeuroImage 44, 489–501 (2009).

67. Buckner, R. L. The cerebellum and cognitive function: 25 years of insight from anatomy and neuroimaging. Neuron 80, 807–815 (2013).

68. King, M., Hernandez-Castillo, C. R., Poldrack, R. A., Ivry, R. B. & Diedrichsen, J. Functional boundaries in the human cerebellum revealed by a multi-domain task battery. Nat. Neurosci. 22, 1371–1378 (2019).

69. Ivry, R. Exploring the role of the cerebellum in sensory anticipation and timing: Commentary on Tesche and Karhu. Hum. Brain Mapp. 9, 115–118 (2000).

70. Roth, M. J., Synofzik, M. & Lindner, A. The Cerebellum Optimizes Perceptual Predictions about External Sensory Events. Curr. Biol. 23, 930–935 (2013).

71. Tanaka, M. et al. Roles of the Cerebellum in Motor Preparation and Prediction of Timing. Neuroscience 462, 220–234 (2021).

72. Palmer, S. J., Li, J., Wang, Z. J. & McKeown, M. J. Joint amplitude and connectivity compensatory mechanisms in Parkinson’s disease. Neuroscience 166, 1110–1118 (2010).

73. Wu, T. & Hallett, M. The cerebellum in Parkinson’s disease. Brain 136, 696–709 (2013).

74. Yu, H., Sternad, D., Corcos, D. M. & Vaillancourt, D. E. Role of hyperactive cerebellum and motor cortex in Parkinson’s disease. NeuroImage 35, 222–233 (2007).

75. Engel, A. K. & Fries, P. Beta-band oscillations--signalling the status quo? Curr. Opin. Neurobiol. 20, 156–165 (2010).

76. Jones, C. R. G. & Jahanshahi, M. Contributions of the Basal Ganglia to Temporal Processing: Evidence from Parkinson’s Disease. Timing Time Percept. 2, 87–127 (2014).

77. Rao, S. M., Mayer, A. R. & Harrington, D. L. The evolution of brain activation during temporal processing. Nat. Neurosci. 4, 317–323 (2001).

78. Kish, S. J., Shannak, K. & Hornykiewicz, O. Uneven pattern of dopamine loss in the striatum of patients with idiopathic Parkinson’s disease. Pathophysiologic and clinical implications. N. Engl. J. Med. 318, 876–880 (1988).

79. Macdonald, P. A. & Monchi, O. Differential effects of dopaminergic therapies on dorsal and ventral striatum in Parkinson’s disease: implications for cognitive function. Park. Dis. 2011, 572743 (2011).

80. Cools, R., Barker, R. A., Sahakian, B. J. & Robbins, T. W. Enhanced or Impaired Cognitive Function in Parkinson’s Disease as a Function of Dopaminergic Medication and Task Demands. Cereb. Cortex 11, 1136–1143 (2001).

81. Kordower, J. H. et al. Disease duration and the integrity of the nigrostriatal system in Parkinson’s disease. Brain J. Neurol. 136, 2419–2431 (2013).

82. Breska, A. & Ivry, R. B. Double dissociation of single-interval and rhythmic temporal prediction in cerebellar degeneration and Parkinson’s disease. Proc. Natl. Acad. Sci. 115, 12283–12288 (2018).

83. Nozaradan, S., Schwartze, M., Obermeier, C. & Kotz, S. A. Specific contributions of basal ganglia and cerebellum to the neural tracking of rhythm. Cortex 95, 156–168 (2017).

84. Teki, S., Grube, M. & Griffiths, T. D. A Unified Model of Time Perception Accounts for Duration-Based and Beat-Based Timing Mechanisms. Front. Integr. Neurosci. 5, (2012).

85. Kühn, A. A., Kupsch, A., Schneider, G.-H. & Brown, P. Reduction in subthalamic 8–35 Hz oscillatory activity correlates with clinical improvement in Parkinson’s disease. Eur. J. Neurosci. 23, 1956–1960 (2006).

86. Bastos, A. M. & Schoffelen, J.-M. A Tutorial Review of Functional Connectivity Analysis Methods and Their Interpretational Pitfalls. Front. Syst. Neurosci. 9, 175 (2015).

87. Tinkhauser, G. et al. Beta burst dynamics in Parkinson’s disease OFF and ON dopaminergic medication Europe PMC Funders Group. Brain 140, 2968–298110 (2017).

88. Schlerf, J. E., Galea, J. M., Spampinato, D. & Celnik, P. A. Laterality Differences in Cerebellar– Motor Cortex Connectivity. Cereb. Cortex 25, 1827–1834 (2015).

89. Wu, T., Hou, Y., Hallett, M., Zhang, J. & Chan, P. Lateralization of brain activity pattern during unilateral movement in Parkinson’s disease. Hum. Brain Mapp. 36, 1878–1891 (2015).

