## Supplementary Information for "Somatosensory rhythms and cerebellar-basal ganglia beta-band dynamics in Parkinson’s disease"

*Results*

A complementary analysis using z-scored responses was performed to evaluate the robustness of the ratio-based approach. The resulting cerebellar and basal ganglia interaction effects were highly similar to those obtained using the original normalization procedure.

For the cerebellum, the reported analysis revealed significant effects from –29 ms to –16 ms, peaking at –22 ms (d = –0.940), whereas the z-scored analysis revealed significant effects from –38 ms to –9 ms, peaking at –24 ms (d = –1.20).

For the basal ganglia, the reported analysis revealed significant effects from –26 ms to –6 ms, peaking at –15 ms (d = 1.09), whereas the z-scored analysis revealed significant effects from –21 ms to –12 ms, peaking at –16 ms (d = 0.953).

Thus, the principal findings were robust to the normalization approach. All analyses were conducted using threshold-free cluster enhancement (TFCE).

***Figure S1. Somatosensory responses*** *for SI (53 ms) and SII (132 ms); (**a**) Controls, n=25; **(b)** Parkinson's disease, n=24. Signal Space Projection (first four SSPs) was applied to reduce environmental noise.*

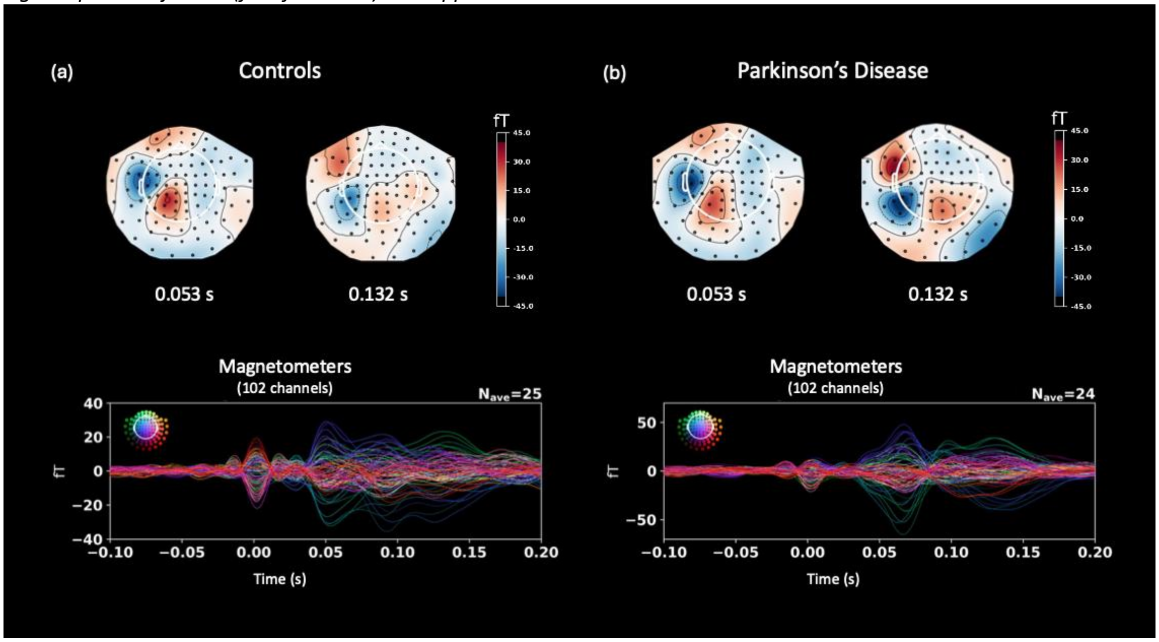

***Figure S2***. *Distribution of prescribed medication for Parkinson's disease symptoms. L-DOPA, levodopa; D-AGONIST, dopamine agonist; MAOI, monoamine oxidase inhibitor; COMT-I; Catechol-O-methyltransferase inhibitor.*

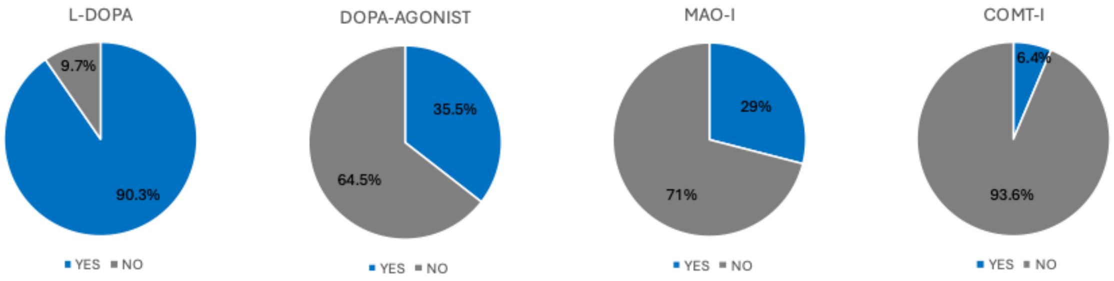

### Table S1. Detailed demographics.

| ID | Group | Age | Sex | UPDRS-III | H & Y | MDI | MoCA | PD dx | L-DOPA | DOPA-AGONIST | MAO-I | COMT-I | LEDD | Reason for exclusion |
|---|---|---|---|---|---|---|---|---|---|---|---|---|---|---|
| 2 | PD | 63 | M | 22 | 1 | 6 | 30 | 1 | YES | NO | YES | NO | 500 | |
| 3 | control | 58 | F | 0 | 0 | 3 | 30 | 0 | NO | NO | NO | NO | 0 | |
| 4 | PD | 63 | M | 25 | 1 | 8 | 30 | 3 | NO | YES | NO | NO | 320 | |
| 5 | control | 56 | M | 0 | 0 | 2 | 28 | 0 | NO | NO | NO | NO | 0 | |
| 6 | PD | 69 | F | 27 | 1 | 6 | 30 | 1 | YES | NO | YES | NO | 500 | |
| 7 | PD | 75 | M | 39 | 2 | 0 | 30 | 15 | YES | YES | YES | NO | 772 | |
| 8 | PD | 55 | M | 28 | 1 | 8 | 29 | 9 | YES | NO | NO | NO | 400 | left-handed |
| 9 | PD | 66 | M | 39 | 2 | 5 | 29 | 10 | YES | NO | YES | NO | 407 | |
| 10 | PD | 66 | M | 38 | 2 | 5 | 27 | 5 | YES | YES | YES | YES | 407 | |
| 11 | PD | 70 | M | 40 | 2 | 5 | 27 | 12 | YES | NO | NO | NO | 470 | |
| 12 | PD | 59 | F | 23 | 1 | 3 | 29 | 10 | YES | YES | NO | NO | 652 | |
| 13 | PD | 70 | F | 31 | 2 | 4 | 28 | 12 | YES | NO | NO | NO | 800 | |
| 14 | PD | 45 | M | 38 | 2 | 10 | 28 | 3 | NO | YES | NO | NO | 320 | left-handed |
| 15 | PD | 69 | M | 38 | 2 | 15 | 27 | 6 | YES | YES | YES | NO | 862 | |
| 16 | control | 66 | F | 0 | 0 | 4 | 30 | 0 | NO | NO | NO | NO | 0 | |
| 17 | control | 39 | F | 0 | 0 | 3 | 29 | 0 | NO | NO | NO | NO | 0 | |
| 18 | PD | 76 | F | 24 | 1 | 11 | 30 | 2 | YES | NO | NO | NO | 400 | |
| 19 | control | 67 | M | 0 | 0 | 7 | 29 | 0 | NO | NO | NO | NO | 0 | left-handed |
| 20 | control | 59 | F | 0 | 0 | 1 | 27 | 0 | NO | NO | NO | NO | 0 | |
| 21 | PD | 74 | M | 39 | 2 | 10 | 29 | 17 | YES | YES | YES | YES | 1014 | |
| 22 | control | 55 | M | 0 | 0 | 2 | 27 | 0 | NO | NO | NO | NO | 0 | |
| 23 | control | 57 | M | 0 | 0 | 4 | 30 | 0 | NO | NO | NO | NO | 0 | |
| 24 | PD | 67 | M | 35 | 2 | 4 | 29 | 2 | YES | YES | YES | NO | 235 | metal plate |
| 25 | PD | 66 | M | 26 | 2 | 3 | 28 | 1 | YES | NO | NO | NO | 500 | metal plate |
| 26 | control | 71 | F | 0 | 0 | 2 | 28 | 0 | NO | NO | NO | NO | 0 | |
| 27 | PD | 48 | M | 13 | 1 | 11 | 27 | 3 | NO | YES | YES | NO | 270 | |
| 28 | PD | 47 | M | 26 | 2 | 15 | 27 | 3 | YES | NO | NO | NO | 400 | |
| 29 | control | 51 | F | 0 | 0 | 5 | 30 | 0 | NO | NO | NO | NO | 0 | |
| 30 | control | 61 | F | 0 | 0 | 13 | 28 | 0 | NO | NO | NO | NO | 0 | |
| 31 | control | 42 | M | 0 | 0 | 7 | 28 | 0 | NO | NO | NO | NO | 0 | |
| 32 | control | 43 | M | 0 | 0 | 3 | 29 | 0 | NO | NO | NO | NO | 0 | |
| 33 | control | 65 | F | 0 | 0 | 0 | 30 | 0 | NO | NO | NO | NO | 0 | |
| 34 | control | 53 | M | 0 | 0 | 10 | 29 | 0 | NO | NO | NO | NO | 0 | |
| 35 | PD | 64 | F | 23 | 2 | 7 | 26 | 5 | YES | YES | NO | NO | 520 | |
| 36 | PD | 67 | F | 37 | 2 | 12 | 28 | 1 | YES | NO | NO | NO | 500 | cerebellar atrophy |
| 37 | control | 52 | F | 0 | 0 | 2 | 29 | 0 | NO | NO | NO | NO | 0 | |
| 38 | control | 61 | F | 0 | 0 | 19 | 27 | 0 | NO | NO | NO | NO | 0 | |
| 39 | control | 69 | M | 0 | 0 | 1 | 26 | 0 | NO | NO | NO | NO | 0 | |
| 40 | PD | 77 | M | 41 | 2 | 17 | 28 | 4 | YES | NO | NO | NO | 520 | metal plate |
| 41 | control | 72 | F | 0 | 0 | 3 | 28 | 0 | NO | NO | NO | NO | 0 | |
| 42 | PD | 57 | F | 30 | 2 | 8 | 30 | 7 | YES | NO | NO | NO | 500 | |
| 43 | PD | 71 | M | 47 | 2 | 10 | 27 | 14 | YES | NO | NO | NO | 1000 | |
| 44 | PD | 73 | M | 31 | 2 | 13 | 30 | 5 | YES | NO | NO | NO | 400 | |
| 45 | control | 69 | F | 0 | 0 | 2 | 26 | 0 | NO | NO | NO | NO | 0 | |
| 46 | PD | 73 | F | 34 | 2 | 8 | 30 | 8 | YES | NO | NO | NO | 600 | |
| 47 | PD | 68 | M | 29 | 2 | 8 | 27 | 2 | YES | NO | NO | NO | 400 | |
| 48 | control | 63 | F | 0 | 0 | 17 | 28 | 0 | NO | NO | NO | NO | 0 | |
| 49 | PD | 57 | F | 43 | 2 | 9 | 29 | 2 | YES | NO | NO | NO | 400 | |
| 50 | PD | 72 | M | 26 | 2 | 15 | 30 | 4 | YES | NO | NO | NO | 400 | |
| 51 | PD | 73 | M | 25 | 2 | 4 | 27 | 4 | YES | NO | NO | NO | 400 | |
| 52 | control | 65 | M | 0 | 0 | 0 | 27 | 0 | NO | NO | NO | NO | 0 | |
| 53 | control | 73 | F | 0 | 0 | 3 | 29 | 0 | NO | NO | NO | NO | 0 | |
| 54 | PD | | | 43 | 2 | 5 | 28 | 2 | YES | NO | NO | NO | 312.5 | |
| 55 | control | | | 0 | 0 | 5 | 30 | 0 | NO | NO | NO | NO | 0 | |
| 56 | control | | | 0 | 0 | 0 | 29 | 0 | NO | NO | NO | NO | 0 | |
| 57 | control | | | 0 | 0 | 4 | 29 | 0 | NO | NO | NO | NO | 0 | |

PD, Parkinson's disease; MDI, Major Depression Inventory; MoCA, Montreal Cognitive Assessment; PD dx, years since PD diagnosis; UPDRS-III, Motor Examination of the Movement Disorder Society Unified Parkinson's Disease Rating Scale; H & Y, Hoehn and Yahr scale; L-DOPA, levodopa; DOPA-AGONIST, dopamine agonist; MAO-I, monoamine oxidase inhibitor; COMT-I, Catechol-O-methyltransferase inhibitor; LEDD, levodopa equivalent daily dose.

### Table S2. Movement Disorder Society (MDS) Unified Parkinson's Disease Rating Scale (UPDRS) – Part III.

| Item/ID | 1 | 2 | 3 | 4 | 5 | 6 | 7 | 8 | 9 | 10 | 11 | 12 | 13 | 14 | 15 | 16 | 17 | 18 | 19 | 20 | 21 | 22 | 23 | 24 | 25 |
|---|---|---|---|---|---|---|---|---|---|---|---|---|---|---|---|---|---|---|---|---|---|---|---|---|---|
| **Year Dx** | 2019 | 2022 | NA | 2019 | NA | 2022 | 2008 | 2014 | 2013 | 2017 | 2011 | 2013 | 2011 | 2020 | 2017 | NA | NA | 2021 | NA | NA | 2006 | NA | NA | 2021 | 2022 |
| **Most affected side** | right | right | NA | left | NA | right | right | right | right | left | right | left | left | left | left | NA | NA | left | NA | NA | left | NA | NA | right | left |
| **Minutes since last levodopa** | 45 | 0 | NA | NA | NA | 90 | 150 | 100 | 0 | 0 | 30 | 90 | NA | 100 | 30 | NA | NA | 30 | NA | NA | 90 | NA | NA | 180 | 240 |
| 3.1 Speech | 0 | 1 | NA | 1 | NA | 0 | 1 | 2 | 2 | 2 | 2 | 1 | 1 | 1 | 1 | NA | NA | 0 | NA | NA | 1 | NA | NA | 0 | 1 |
| 3.2 Facial expression | 1 | 1 | NA | 1 | NA | 0 | 1 | 1 | 1 | 2 | 2 | 1 | 1 | 2 | 2 | NA | NA | 1 | NA | NA | 2 | NA | NA | 1 | 1 |
| 3.3a Rigidity– Neck | 1 | 0 | NA | 1 | NA | 1 | 2 | 1 | 2 | 1 | 2 | 1 | 1 | 1 | 1 | NA | NA | 0 | NA | NA | 1 | NA | NA | 1 | 1 |
| 3.3b Rigidity– RUE | 2 | 2 | NA | 1 | NA | 3 | 3 | 2 | 2 | 1 | 2 | 0 | 1 | 1 | 1 | NA | NA | 1 | NA | NA | 1 | NA | NA | 1 | 1 |
| 3.3c Rigidity– LUE | 1 | 1 | NA | 1 | NA | 0 | 1 | 1 | 1 | 2 | 2 | 1 | 1 | 1 | 2 | NA | NA | 1 | NA | NA | 1 | NA | NA | 1 | 2 |
| 3.3d Rigidity– RLE | 1 | 0 | NA | 1 | NA | 2 | 2 | 2 | 2 | 1 | 3 | 0 | 2 | 2 | 1 | NA | NA | 1 | NA | NA | 1 | NA | NA | 2 | 1 |
| 3.3e Rigidity– LLE | 1 | 0 | NA | 1 | NA | 0 | 1 | 1 | 1 | 2 | 1 | 1 | 3 | 2 | 2 | NA | NA | 1 | NA | NA | 2 | NA | NA | 3 | 2 |
| 3.4a Finger tapping– Right hand | 2 | 2 | NA | 1 | NA | 3 | 2 | 2 | 3 | 1 | 2 | 1 | 1 | 1 | 1 | NA | NA | 1 | NA | NA | 1 | NA | NA | 2 | 1 |
| 3.4b Finger tapping– Left hand | 1 | 1 | NA | 2 | NA | 0 | 1 | 1 | 1 | 3 | 1 | 2 | 2 | 2 | 2 | NA | NA | 2 | NA | NA | 2 | NA | NA | 2 | 2 |
| 3.5a Hand movements– Right hand | 1 | 2 | NA | 1 | NA | 3 | 3 | 2 | 3 | 1 | 2 | 1 | 1 | 1 | 1 | NA | NA | 1 | NA | NA | 1 | NA | NA | 1 | 1 |
| 3.5b Hand movements– Left hand | 1 | 1 | NA | 2 | NA | 0 | 1 | 1 | 1 | 3 | 2 | 2 | 2 | 2 | 2 | NA | NA | 2 | NA | NA | 3 | NA | NA | 1 | 2 |
| 3.6a Pronation-supination – Right hand | 2 | 2 | NA | 1 | NA | 3 | 3 | 2 | 3 | 1 | 2 | 1 | 1 | 1 | 1 | NA | NA | 1 | NA | NA | 1 | NA | NA | 2 | 1 |
| 3.6b Pronation-supination – Left hand | 1 | 1 | NA | 2 | NA | 0 | 1 | 1 | 1 | 3 | 2 | 2 | 2 | 2 | 1 | NA | NA | 2 | NA | NA | 3 | NA | NA | 1 | 2 |
| 3.7a Toe tapping– Right foot | 0 | 1 | NA | 0 | NA | 1 | 0 | 0 | 0 | 0 | 0 | 0 | 0 | 0 | 0 | NA | NA | 0 | NA | NA | 0 | NA | NA | 2 | 0 |
| 3.7b Toe tapping– Left foot | 0 | 1 | NA | 0 | NA | 0 | 0 | 0 | 0 | 0 | 0 | 1 | 0 | 0 | 0 | NA | NA | 0 | NA | NA | 0 | NA | NA | 1 | 0 |
| 3.8a Leg agility– Right leg | 0 | 0 | NA | 0 | NA | 1 | 2 | 3 | 2 | 1 | 1 | 1 | 1 | 1 | 1 | NA | NA | 0 | NA | NA | 1 | NA | NA | 3 | 0 |
| 3.8b Leg agility– Left leg | 0 | 0 | NA | 0 | NA | 0 | 1 | 2 | 1 | 3 | 0 | 2 | 2 | 2 | 2 | NA | NA | 1 | NA | NA | 1 | NA | NA | 2 | 1 |
| 3.9 Arising from chair | 0 | 0 | NA | 1 | NA | 0 | 1 | 0 | 0 | 0 | 0 | 0 | 0 | 0 | 1 | NA | NA | 0 | NA | NA | 0 | NA | NA | 0 | 0 |
| 3.10 Gait | 0 | 0 | NA | 0 | NA | 1 | 1 | 0 | 2 | 1 | 1 | 0 | 1 | 0 | 2 | NA | NA | 2 | NA | NA | 0 | NA | NA | 0 | 0 |
| 3.11 Freezing of gait | 0 | 0 | NA | 0 | NA | 0 | 0 | 0 | 0 | 0 | 0 | 0 | 0 | 0 | 0 | NA | NA | 0 | NA | NA | 0 | NA | NA | 0 | 0 |
| 3.12 Postural stability | 0 | 0 | NA | 0 | NA | 0 | 1 | 1 | 1 | 1 | 1 | 0 | 1 | 0 | 1 | NA | NA | 1 | NA | NA | 0 | NA | NA | 0 | 0 |
| 3.13 Posture | 0 | 0 | NA | 0 | NA | 0 | 2 | 0 | 1 | 1 | 1 | 0 | 1 | 0 | 2 | NA | NA | 1 | NA | NA | 2 | NA | NA | 0 | 0 |
| 3.14 Global spontaneity of movement | 1 | 1 | NA | 1 | NA | 1 | 2 | 1 | 2 | 2 | 2 | 1 | 1 | 1 | 2 | NA | NA | 1 | NA | NA | 2 | NA | NA | 1 | 1 |
| 3.15a Postural tremor– Right hand | 1 | 1 | NA | 1 | NA | 2 | 2 | 1 | 1 | 1 | 2 | 1 | 1 | 3 | 2 | NA | NA | 0 | NA | NA | 2 | NA | NA | 1 | 1 |
| 3.15b Postural tremor– Left hand | 0 | 0 | NA | 1 | NA | 0 | 0 | 0 | 0 | 1 | 1 | 1 | 2 | 3 | 1 | NA | NA | 1 | NA | NA | 3 | NA | NA | 1 | 2 |
| 3.16a Kinetic tremor– Right hand | 1 | 1 | NA | 1 | NA | 2 | 1 | 0 | 3 | 1 | 2 | 0 | 0 | 1 | 2 | NA | NA | 0 | NA | NA | 2 | NA | NA | 1 | 0 |
| 3.16b Kinetic tremor– Left hand | 0 | 0 | NA | 1 | NA | 0 | 1 | 0 | 1 | 1 | 1 | 1 | 0 | 1 | 2 | NA | NA | 1 | NA | NA | 2 | NA | NA | 1 | 0 |
| 3.17a Rest tremor amplitude– RUE | 1 | 1 | NA | 0 | NA | 2 | 0 | 0 | 0 | 0 | 0 | 0 | 0 | 2 | 0 | NA | NA | 0 | NA | NA | 0 | NA | NA | 1 | 0 |
| 3.17b Rest tremor amplitude– LUE | 0 | 0 | NA | 1 | NA | 0 | 0 | 0 | 0 | 0 | 0 | 0 | 0 | 2 | 0 | NA | NA | 0 | NA | NA | 1 | NA | NA | 0 | 0 |
| 3.17c Rest tremor amplitude– RLE | 0 | 0 | NA | 0 | NA | 0 | 0 | 0 | 0 | 0 | 0 | 0 | 0 | 0 | 0 | NA | NA | 0 | NA | NA | 0 | NA | NA | 0 | 0 |
| 3.17d Rest tremor amplitude– LLE | 0 | 0 | NA | 0 | NA | 0 | 0 | 0 | 0 | 0 | 0 | 0 | 0 | 0 | 0 | NA | NA | 0 | NA | NA | 0 | NA | NA | 0 | 0 |
| 3.17e Rest tremor amplitude– Lip/jaw | 0 | 0 | NA | 0 | NA | 0 | 0 | 0 | 0 | 0 | 0 | 0 | 0 | 0 | 0 | NA | NA | 0 | NA | NA | 0 | NA | NA | 0 | 0 |
| 3.18 Constancy of rest tremor | 0 | 1 | NA | 1 | NA | 1 | 1 | 0 | 0 | 0 | 1 | 0 | 0 | 1 | 0 | NA | NA | 1 | NA | NA | 1 | NA | NA | 1 | 1 |
| Hoehn and Yahr stage (0-5) | 1 | 1 | NA | 1 | NA | 1 | 2 | 1 | 2 | 2 | 2 | 1 | 2 | 2 | 2 | NA | NA | 1 | NA | NA | 2 | NA | NA | 2 | 2 |
| Were dyskinesias present? y/n | no | no | NA | no | NA | no | no | no | no | no | yes | no | no | no | yes | NA | NA | no | NA | NA | no | NA | NA | no | no |
| Did they interfere with ratings? y/n | no | no | NA | no | NA | no | no | no | no | no | no | no | no | no | no | NA | NA | no | NA | NA | no | NA | NA | no | no |
| **Total Part III Score** | **20** | **22** | **0** | **25** | **0** | **27** | **39** | **28** | **39** | **38** | **40** | **23** | **31** | **38** | **38** | **0** | **0** | **24** | **0** | **0** | **39** | **0** | **0** | **35** | **26** |

| Item/ID | 26 | 27 | 28 | 29 | 30 | 31 | 32 | 33 | 34 | 35 | 36 | 37 | 38 | 39 | 40 | 41 | 42 | 43 | 44 | 45 | 46 | 47 | 48 | 49 | 50 | 51 | 52 | 53 | 54 | 55 | 56 | 57 |
|---|---|---|---|---|---|---|---|---|---|---|---|---|---|---|---|---|---|---|---|---|---|---|---|---|---|---|---|---|---|---|---|---|
| **Year Dx** | NA | 2020 | 2020 | NA | NA | NA | NA | NA | NA | 2018 | 2023 | NA | NA | NA | 2019 | NA | 2016 | 2009 | 2018 | NA | 2015 | 2021 | NA | 2021 | 2019 | 2019 | NA | NA | 2021 | NA | NA | NA |
| **Most affected side** | NA | left | right | NA | NA | NA | NA | NA | NA | left | left | NA | NA | NA | right | NA | left | left | left | NA | right | right | NA | right | right | right | NA | NA | left | NA | NA | NA |
| **Minutes since last levodopa** | NA | 120 | 180 | NA | NA | NA | NA | NA | NA | 90 | 240 | NA | NA | NA | 180 | NA | 105 | 0 | 90 | NA | 15 | 75 | NA | 90 | 180 | 180 | NA | NA | 120 | NA | NA | NA |
| 3.1 Speech | NA | 0 | 0 | NA | NA | NA | NA | NA | NA | 0 | 1 | NA | NA | NA | 2 | NA | 0 | 2 | 0 | NA | 2 | 0 | NA | 0 | 0 | 2 | NA | NA | 1 | NA | NA | NA |
| 3.2 Facial expression | NA | 0 | 2 | NA | NA | NA | NA | NA | NA | 1 | 2 | NA | NA | NA | 2 | NA | 1 | 3 | 1 | NA | 2 | 1 | NA | 1 | 1 | 2 | NA | NA | 2 | NA | NA | NA |
| 3.3a Rigidity– Neck | NA | 0 | 1 | NA | NA | NA | NA | NA | NA | 0 | 2 | NA | NA | NA | 2 | NA | 0 | 2 | 2 | NA | 2 | 1 | NA | 1 | 0 | 0 | NA | NA | 2 | NA | NA | NA |
| 3.3b Rigidity– RUE | NA | 0 | 1 | NA | NA | NA | NA | NA | NA | 1 | 2 | NA | NA | NA | 1 | NA | 2 | 2 | 1 | NA | 2 | 2 | NA | 2 | 1 | 0 | NA | NA | 1 | NA | NA | NA |
| 3.3c Rigidity– LUE | NA | 0 | 1 | NA | NA | NA | NA | NA | NA | 1 | 2 | NA | NA | NA | 1 | NA | 2 | 2 | 1 | NA | 1 | 2 | NA | 1 | 2 | 0 | NA | NA | 1 | NA | NA | NA |
| 3.3d Rigidity– RLE | NA | 0 | 2 | NA | NA | NA | NA | NA | NA | 1 | 1 | NA | NA | NA | 1 | NA | 1 | 2 | 1 | NA | 2 | 1 | NA | 2 | 2 | 1 | NA | NA | 1 | NA | NA | NA |
| 3.3e Rigidity– LLE | NA | 0 | 1 | NA | NA | NA | NA | NA | NA | 1 | 2 | NA | NA | NA | 1 | NA | 1 | 2 | 1 | NA | 1 | 1 | NA | 1 | 1 | 1 | NA | NA | 1 | NA | NA | NA |
| 3.4a Finger tapping– Right hand | NA | 0 | 2 | NA | NA | NA | NA | NA | NA | 1 | 1 | NA | NA | NA | 2 | NA | 0 | 1 | 1 | NA | 2 | 2 | NA | 3 | 1 | 2 | NA | NA | 2 | NA | NA | NA |
| 3.4b Finger tapping– Left hand | NA | 2 | 1 | NA | NA | NA | NA | NA | NA | 2 | 2 | NA | NA | NA | 2 | NA | 2 | 2 | 3 | NA | 1 | 1 | NA | 2 | 1 | 1 | NA | NA | 3 | NA | NA | NA |
| 3.5a Hand movements– Right hand | NA | 0 | 2 | NA | NA | NA | NA | NA | NA | 1 | 1 | NA | NA | NA | 3 | NA | 0 | 1 | 1 | NA | 2 | 2 | NA | 3 | 1 | 2 | NA | NA | 2 | NA | NA | NA |
| 3.5b Hand movements– Left hand | NA | 3 | 1 | NA | NA | NA | NA | NA | NA | 2 | 2 | NA | NA | NA | 3 | NA | 2 | 2 | 3 | NA | 1 | 1 | NA | 2 | 1 | 1 | NA | NA | 3 | NA | NA | NA |
| 3.6a Pronation-supination – Right hand | NA | 0 | 2 | NA | NA | NA | NA | NA | NA | 1 | 1 | NA | NA | NA | 2 | NA | 1 | 1 | 1 | NA | 1 | 1 | NA | 3 | 1 | 2 | NA | NA | 2 | NA | NA | NA |
| 3.6b Pronation-supination – Left hand | NA | 2 | 1 | NA | NA | NA | NA | NA | NA | 2 | 2 | NA | NA | NA | 2 | NA | 2 | 2 | 2 | NA | 0 | 1 | NA | 2 | 1 | 1 | NA | NA | 3 | NA | NA | NA |
| 3.7a Toe tapping– Right foot | NA | 0 | 0 | NA | NA | NA | NA | NA | NA | 0 | 1 | NA | NA | NA | 1 | NA | 0 | 0 | 0 | NA | 1 | 1 | NA | 1 | 0 | 1 | NA | NA | 1 | NA | NA | NA |
| 3.7b Toe tapping– Left foot | NA | 0 | 0 | NA | NA | NA | NA | NA | NA | 1 | 1 | NA | NA | NA | 1 | NA | 1 | 1 | 1 | NA | 0 | 1 | NA | 0 | 0 | 0 | NA | NA | 2 | NA | NA | NA |
| 3.8a Leg agility– Right leg | NA | 1 | 0 | NA | NA | NA | NA | NA | NA | 1 | 1 | NA | NA | NA | 1 | NA | 1 | 1 | 1 | NA | 1 | 1 | NA | 3 | 1 | 1 | NA | NA | 1 | NA | NA | NA |
| 3.8b Leg agility– Left leg | NA | 2 | 0 | NA | NA | NA | NA | NA | NA | 1 | 2 | NA | NA | NA | 1 | NA | 2 | 2 | 3 | NA | 0 | 1 | NA | 2 | 1 | 0 | NA | NA | 2 | NA | NA | NA |
| 3.9 Arising from chair | NA | 0 | 1 | NA | NA | NA | NA | NA | NA | 0 | 0 | NA | NA | NA | 0 | NA | 1 | 3 | 0 | NA | 0 | 0 | NA | 1 | 0 | 0 | NA | NA | 0 | NA | NA | NA |
| 3.10 Gait | NA | 0 | 0 | NA | NA | NA | NA | NA | NA | 1 | 2 | NA | NA | NA | 1 | NA | 0 | 2 | 0 | NA | 1 | 1 | NA | 3 | 1 | 1 | NA | NA | 2 | NA | NA | NA |
| 3.11 Freezing of gait | NA | 0 | 0 | NA | NA | NA | NA | NA | NA | 0 | 1 | NA | NA | NA | 1 | NA | 1 | 1 | 0 | NA | 1 | 1 | NA | 0 | 0 | 0 | NA | NA | 0 | NA | NA | NA |
| 3.12 Postural stability | NA | 0 | 0 | NA | NA | NA | NA | NA | NA | 0 | 0 | NA | NA | NA | 0 | NA | 0 | 0 | 0 | NA | 0 | 0 | NA | 1 | 0 | 1 | NA | NA | 0 | NA | NA | NA |
| 3.13 Posture | NA | 0 | 0 | NA | NA | NA | NA | NA | NA | 1 | 0 | NA | NA | NA | 1 | NA | 0 | 2 | 0 | NA | 1 | 1 | NA | 1 | 1 | 1 | NA | NA | 1 | NA | NA | NA |
| 3.14 Global spontaneity of movement | NA | 0 | 0 | NA | NA | NA | NA | NA | NA | 1 | 1 | NA | NA | NA | 1 | NA | 1 | 2 | 1 | NA | 2 | 2 | NA | 2 | 1 | 1 | NA | NA | 3 | NA | NA | NA |
| 3.15a Postural tremor– Right hand | NA | 0 | 2 | NA | NA | NA | NA | NA | NA | 1 | 1 | NA | NA | NA | 2 | NA | 1 | 1 | 1 | NA | 1 | 1 | NA | 2 | 1 | 1 | NA | NA | 1 | NA | NA | NA |
| 3.15b Postural tremor– Left hand | NA | 1 | 1 | NA | NA | NA | NA | NA | NA | 0 | 2 | NA | NA | NA | 2 | NA | 1 | 2 | 1 | NA | 1 | 1 | NA | 1 | 1 | 0 | NA | NA | 1 | NA | NA | NA |
| 3.16a Kinetic tremor– Right hand | NA | 0 | 2 | NA | NA | NA | NA | NA | NA | 0 | 1 | NA | NA | NA | 1 | NA | 0 | 1 | 1 | NA | 1 | 0 | NA | 0 | 1 | 0 | NA | NA | 1 | NA | NA | NA |
| 3.16b Kinetic tremor– Left hand | NA | 1 | 1 | NA | NA | NA | NA | NA | NA | 0 | 1 | NA | NA | NA | 1 | NA | 2 | 1 | 1 | NA | 1 | 0 | NA | 0 | 1 | 0 | NA | NA | 1 | NA | NA | NA |
| 3.17a Rest tremor amplitude– RUE | NA | 0 | 0 | NA | NA | NA | NA | NA | NA | 0 | 0 | NA | NA | NA | 1 | NA | 0 | 0 | 0 | NA | 1 | 0 | NA | 1 | 1 | 1 | NA | NA | 0 | NA | NA | NA |
| 3.17b Rest tremor amplitude– LUE | NA | 0 | 0 | NA | NA | NA | NA | NA | NA | 0 | 0 | NA | NA | NA | 0 | NA | 2 | 1 | 0 | NA | 0 | 0 | NA | 0 | 0 | 0 | NA | NA | 1 | NA | NA | NA |
| 3.17c Rest tremor amplitude– RLE | NA | 0 | 0 | NA | NA | NA | NA | NA | NA | 0 | 0 | NA | NA | NA | 0 | NA | 0 | 0 | 0 | NA | 0 | 0 | NA | 0 | 0 | 0 | NA | NA | 0 | NA | NA | NA |
| 3.17d Rest tremor amplitude– LLE | NA | 0 | 0 | NA | NA | NA | NA | NA | NA | 0 | 0 | NA | NA | NA | 0 | NA | 0 | 0 | 0 | NA | 0 | 0 | NA | 0 | 0 | 0 | NA | NA | 0 | NA | NA | NA |
| 3.17e Rest tremor amplitude– Lip/jaw | NA | 0 | 0 | NA | NA | NA | NA | NA | NA | 0 | 0 | NA | NA | NA | 0 | NA | 0 | 0 | 0 | NA | 0 | 0 | NA | 0 | 0 | 0 | NA | NA | 0 | NA | NA | NA |
| 3.18 Constancy of rest tremor | NA | 0 | 0 | NA | NA | NA | NA | NA | NA | 0 | 0 | NA | NA | NA | 0 | NA | 1 | 1 | 1 | NA | 1 | 0 | NA | 0 | 1 | 0 | NA | NA | 0 | NA | NA | NA |
| Hoehn and Yahr stage (0-5) | NA | 1 | 2 | NA | NA | NA | NA | NA | NA | 2 | 2 | NA | NA | NA | 2 | NA | 2 | 2 | 2 | NA | 2 | 2 | NA | 2 | 2 | 2 | NA | NA | 2 | NA | NA | NA |
| Were dyskinesias present? y/n | NA | no | yes | NA | NA | NA | NA | NA | NA | no | yes | NA | NA | NA | no | NA | no | yes | no | NA | no | no | NA | no | no | no | NA | NA | no | NA | NA | NA |
| Did they interfere with ratings? y/n | NA | no | no | NA | NA | NA | NA | NA | NA | no | no | NA | NA | NA | no | NA | no | no | no | NA | no | no | NA | no | no | no | NA | NA | no | NA | NA | NA |
| **Total Part III Score** | **0** | **13** | **26** | **0** | **0** | **0** | **0** | **0** | **0** | **23** | **37** | **0** | **0** | **0** | **41** | **0** | **30** | **47** | **31** | **0** | **34** | **29** | **0** | **43** | **26** | **25** | **0** | **0** | **43** | **0** | **0** | **0** |

NA, not applicable (control participant); most affected side was determined from the MDS-UPDRS-III by summing left- and right-sided motor item scores and identifying the side with the higher total score); RUE, right upper extremity; LUE, left upper extremity; RLE, right lower extremity; LLE, left lower extremity.
